# ParSe 2.0: A web application that enables proteome-scale searches for sequences that drive protein-mediated phase separation

**DOI:** 10.1101/2023.06.20.545714

**Authors:** Colorado Wilson, Karen A. Lewis, Nicholas C. Fitzkee, Loren E. Hough, Steven T. Whitten

**Affiliations:** Department of Chemistry and Biochemistry, Texas State University, San Marcos, TX, USA; Department of Chemistry, Mississippi State University, Mississippi State, MS, USA; Department of Physics, University of Colorado Boulder, Boulder, CO, USA; BioFrontiers Institute, University of Colorado Boulder, Boulder, CO, USA

## Abstract

We have developed an algorithm, ParSe, that accurately identifies from the primary sequence those protein regions likely to exhibit physiological phase separation behavior. Originally, ParSe was designed to test the hypothesis that, for flexible proteins, phase separation potential is correlated to hydrodynamic size. While our results were consistent with that idea, we also found that many different descriptors could successfully differentiate between three classes of protein regions: folded, intrinsically disordered, and phase-separating intrinsically disordered. Consequently, numerous combinations of amino acid property scales can be used to make robust predictions of protein phase separation. Built from that finding, ParSe 2.0 uses an optimal set of property scales to predict domain-level organization and compute a sequence-based prediction of phase separation potential. The algorithm is fast enough to scan the whole of the human proteome in minutes on a single computer and is equally or more accurate than other published predictors in identifying proteins and regions within proteins that drive phase separation. Here, we describe a web application for ParSe 2.0 that may be accessed through a browser by visiting https://stevewhitten.github.io/Parse_v2_FASTA to quickly identify phase-separating proteins within large sequence sets, or by visiting https://stevewhitten.github.io/Parse_v2_web to evaluate individual protein sequences.

## Introduction

Protein-mediated macromolecular phase separation, through which membrane-free coacervates form spontaneously from the cellular milieu (1–4), is increasingly recognized as an important organizing phenomenon in cells (5–7). By forming specific compartments and micro-environments, protein-mediated macromolecular phase separation, or, more generally, protein phase separation, exerts control over the biochemical reactivity within cells (8–10). Biological coacervates can form in response to environmental stress (11, 12), at specific points in the cell cycle (13, 14), or exist constitutively (3), and have been found to facilitate key cellular processes, including transcription, translation, RNA processing, DNA damage repair, signaling, and metabolism (7, 15–19). Moreover, dysregulation of protein phase separation has been associated with several human diseases (20–22), for example neurodegeneration (4, 23) and cancer (24).

While phase separation can be driven by multivalent interactions between many types of protein domains, including ordered domains (24, 25), many proteins that drive phase separation have intrinsically disordered regions (IDRs) that are necessary and sufficient for phase separation to occur (5, 26–29). Accurate identification of IDRs that drive phase separation is important for testing the underlying mechanisms of phase separation, identifying biological processes that rely on phase separation, and designing sequences that modulate phase separation. To this end, we created the ParSe algorithm (<u>Par</u>tition <u>Se</u>quence; voiced as “parse”). ParSe identifies phase-separating (PS) IDRs starting from predictions of hydrodynamic size (30). The correlation between PS IDR potential and hydrodynamic size assumes that the same forces that drive compaction in monomeric proteins also drive protein phase separation (31–34). Our results were consistent with that idea (30). However, we also found robust property differences between folded, ID, and PS ID protein regions (35). In ParSe 2.0, an optimal set of property scales allows facile predictions of domain-level structure and provides a simple, quantitative metric for the sequence-calculated phase separation potential. Notably, the ParSe-computed PS potential can be modified to account for interactions between amino acids and trained to reproduce effects of mutations on phase separation behavior (35).

A benefit to using ParSe 2.0, compared to the many other available protein phase separation predictors (27, 36–39), is that it can be broadly applied for analyses on very large scales, even to entire proteomes. The algorithm is computationally simple and fast enough to scan tens of thousands of sequences in minutes using a single computer. Moreover, its algorithmic simplicity does not diminish its accuracy; we have found ParSe 2.0 to be as, or more, accurate than other published predictors in identifying proteins and regions within proteins that drive phase separation (35). We have created a web application that enables researchers to utilize ParSe 2.0 for proteome-scale searches for sequences that drive protein phase separation. Herein, we describe the ParSe 2.0 algorithm and show how this application can quickly search large sets of sequences for proteins and the regions within proteins that are predicted to drive phase separation. Also, we show how the ParSe-computed PS potential can be used to predict mutant phase separation behavior, finding that it reasonably reproduced a newly published dataset (40) of mutation effects on the saturation concentration (*c*_*sat*_) associated with protein phase separation.

## Results

The ParSe 2.0 algorithm performs three tasks to resolve the regions within a protein that are ID, and which subset of those are likely to drive phase separation in a biological context. The tasks are:

I. Calculate local properties in the sequence using an optimal set of property scales.
II. Determine where local properties match the folded, ID, or PS ID classes.
III. Identify regions of uniform class to predict domain-level organization.

### Calculate local sequence properties

The algorithm predicts the modular organization within a protein from its regional variations in intrinsic sequence properties. ParSe 2.0 continuously determines the average properties within a 25-residue segment, or window, that advances through the whole sequence, as shown in Figure 1A. This approach avoids averaging properties between distant regions that may have different characteristics.

**Figure 1.**
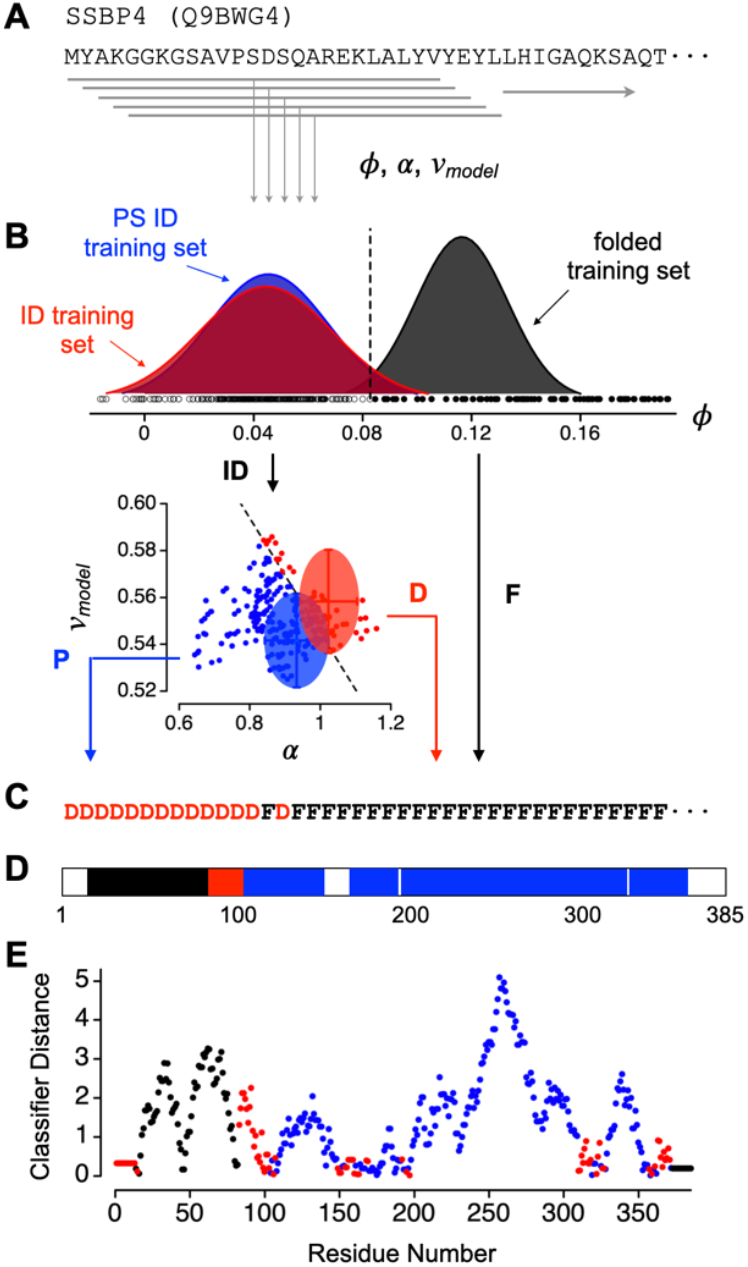
The ParSe 2.0 algorithm. **A)** A sliding window approach is used to identify regions within a protein that match the folded, ID, and PS ID classes. Hydrophobicity (*ϕ*), α-helix propensity (α), and *v*_*model*_ are calculated for each contiguous stretch of 25-residues, or “window”, in the primary sequence. **B)** Each window is assigned a label, F, P, or D, depending on the values of *ϕ*, α, and *v*_*model*_. Small circles show the values calculated for the 25-residue windows in the human SSBP4 sequence; *ϕ* in the top plot, α and *v*_*model*_ in the bottom plot, and compared to the distributions calculated in the folded (black), ID (red), and PS ID training sets (blue). **C)** Window labels are assigned to the central residue of the window. Terminal residues are assigned labels according to the first and last windows. **D)** Contiguous regions of at least 20 residues that are 90% of only one label P, D, or F are colored blue, red, or black, respectively, to represent predicted PS, ID, or folded regions. **E)** Classifier distance of each window, assigned to the central residue of the window and colored according to its label P (blue), D (red), or F (black).

Several properties are calculated from the sequence in each 25-residue segment: 1) the average hydrophobicity, *ϕ*, using an amino acid scale from Bastolla *et al*. that was derived from contact matrices of globular protein structures (41), 2) the average intrinsic propensity for α-helix, α, using an amino acid scale from Tanaka and Scheraga calculated from x-ray data on native proteins (42), and 3) *v*_*model*_, which is a sequence-based model of the polymer scaling exponent (30) that is based on hydrodynamic size and originally was developed from polymer theories to extract information on the balance of self and solvent interactions in long homopolymers (43). Experimental *v* has been used widely as a measure of chain compaction in biological proteins (44–51). The results from this algorithm are relatively independent of the size of the segment or window used (30).

### Match local sequence properties to protein class

Previously, we established that the three classes of proteins regions, for folded, ID, and PS ID, exhibit robust property differences (35). This was shown using curated datasets of folded, ID, and PS ID sequences to examine how broadly existing amino acid property scales can be used to distinguish between the three classes of protein regions. We found that ∼95% of the 566 scales of amino acid properties in the Amino Acid Index Database (52), a curated set of numerical indices representing various physicochemical and biochemical properties of the amino acids, produced statistically significant differences between the means of the folded and ID sets. The largest statistical separation in the means, determined by *t*-test (53), was obtained when using the Bastolla scale for hydrophobicity, *ϕ*. Based on that finding, ParSe 2.0 uses *ϕ* to identify those 25-residue windows in a sequence that are likely to map to a folded protein region (Figure 1B).

Similarly, the largest statistical separation in the means when comparing the ID and PS ID sequence sets was obtained when using the Tanaka and Scheraga propensity scale, α (Figure 1B); however, there was considerable overlap in the performance of different predictors (35). Principal component analysis (PCA) of the variance in the combined sequence sets demonstrated that the variance arising from *v*_*model*_ was orthogonal to the variance from α, and thus α and *v*_*model*_ could be combined without significant redundancy when comparing protein sequences (35). PCA also was used to reduce the dimensionality in the dataset (54), finding that most of the variability within the sequence sets measured by high-performing scales can be captured by 2 to 3 parameters (35). Based on that finding, ParSe 2.0 first uses *ϕ* to identify ID regions (PS ID or ID) as compared to folded regions. Then, it uses α and *v*_*model*_ to identify the 25-residue windows that are likely to show phase separation behavior.

Window labels are used by ParSe 2.0 to record the results of this decision tree. Windows that match the folded class in *ϕ* are labeled F; all others are labeled P or D. D is assigned to windows with high α and high *v*_*model*_ (matching the ID class), while P is assigned to windows with low α and low *v*_*model*_ (matching the PS ID class). The P/D boundary was determined by the line that bisects the overlapping distributions of α and *v*_*model*_ in the ID and PS ID training sets. Next, the window label is assigned to the central residue in the window (Figure 1C). N- and C-terminal residues not belonging to a central window position are assigned the label of the central residue in the first and last window, respectively, of the whole sequence.

### Identify regions of uniform protein class

Protein regions predicted by ParSe 2.0 to be folded, ID, or PS ID are determined by finding contiguous residue positions of length ≥20 that are ≥90% of only one label F, D, or P, respectively. When an overlap occurs between adjacent predicted regions, owing to the up to 10% label mixing allowed, this overlap is split evenly between the two adjacent regions. Figure 1D shows the application of this ruleset to human single-stranded DNA-binding protein 4 (SSBP4), which is not reported to phase separate in the current literature. For SSBP4, protein regions predicted by ParSe 2.0 to be folded, ID, or PS ID have been colored black, red, or blue, respectively; white corresponds to regions with a mixture of F, D, or P labels.

The *classifier distance* was developed to assess confidence in the P, D, and F label assignments (35). Here, the algorithm calculates the linear distance of a window into its classifier sector, relative to the cutoff boundary and normalized by the distance separating the boundary and the training set mean. For example, the classifier distance for a P-labeled window would be the shortest distance of the window to the P/D boundary divided by the shortest distance of the P/D boundary to the mean of the PS ID training set (see Figure 1B). Thus, for a P-labeled window, values greater than 1 in the classifier distance indicate a window located at a distance further from the P/D boundary than that of the PS ID set mean, whereas values less than 1 indicate a window closer to the cutoff boundary than the PS ID set mean and, as such, possibly with some uncertainty for its classifier label (i.e., a classifier distance that resides within the overlap in the ID and PS ID set distributions). For D- and F-labeled windows, identically structured calculations are performed using cutoff boundaries and training set means, for D-labeled windows in the D sector of the α versus *v*_*model*_ plot and for F-labeled windows in the high *ϕ* region. Position-specific classifier distances calculated from the SSBP4 sequence are shown in Figure 1E.

### Proteome-scale searches using ParSe 2.0

One advantage of ParSe 2.0 is that it is very fast and can be applied to very large datasets. To facilitate searches on a proteomic scale, we have developed the ParSe 2.0 algorithm into a web application that takes a user-supplied input in FASTA format. This application may be accessed through a browser by visiting https://stevewhitten.github.io/Parse_v2_FASTA. The required computation time increases linearly with the number of sequences in the input FASTA file. All calculations are performed on the user’s local system through the JavaScript interface. We have found that the application can process, on average, ∼14,000 primary sequences per minute using a standard desktop computer. Figure 2 shows the computational expense in minutes for FASTA files containing various-sized sequence sets. The largest proteome used for this figure is the human proteome with splice variants obtained from UniProt (55) and representing 75,776 primary sequences. The second largest proteome in the figure, also obtained from UniProt, is the human proteome represented by one sequence per gene and containing 20,594 primary sequences. The computation rate for the ParSe 2.0 web application is compared in the figure to the rates we measured for other available tools used to predict phase separation behavior (27, 37–39). ParSe 2.0 is compared in more detail to other predictors below.

**Figure 2.**
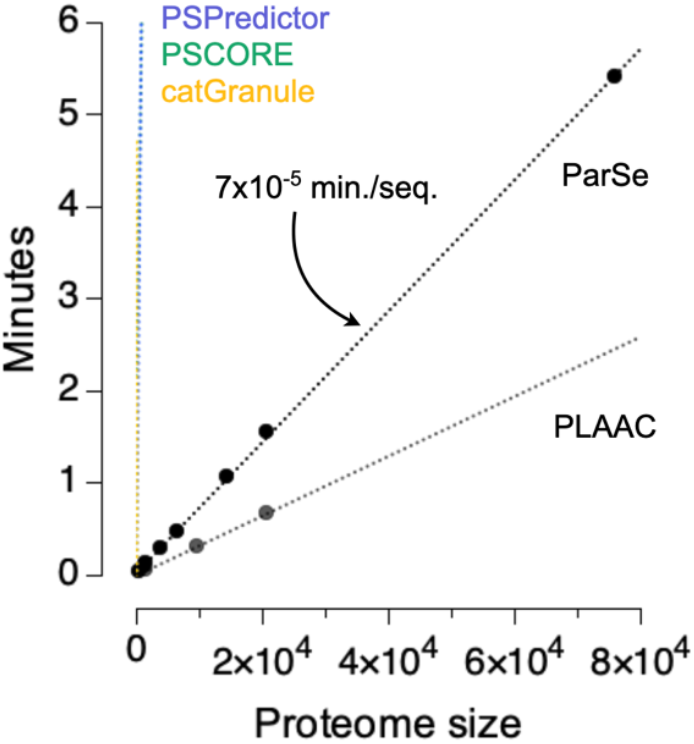
The ParSe 2.0 web application evaluates sequences at a rate of 14,000 primary sequences per minute. Proteome size is the number of primary sequences. The arrow highlights the linear regression slope of the ParSe trend. PSPredictor, PScore, and catGranule are shown in blue, green, and orange, respectively, and overlay in the plot owing to the scale used. The average computation rates for PSPredictor, PScore, catGranule, and PLAAC were 99, 136, 34, and 31,600 primary sequences per minute, respectively. Computation times were determined by using the appropriate web application or python script (see Methods) on a 2015 model iMac.

Upon completion, the application outputs datasets that allow the user to quickly identify those proteins within the input file that have regions predicted to drive phase separation behavior. We demonstrate how to use this application and read its output by example below. We also made a second application (accessed at https://stevewhitten.github.io/Parse_v2_web) that can be used to evaluate individual protein sequences, which produces output in the format shown in Figures 1C-E when provided a single, primary sequence as input.

### Characteristics of biological proteins that drive physiological phase separation

ParSe 2.0 was designed to identify PS IDRs from the primary sequence (35). First, to demonstrate the output expected from proteins that phase separate, we tested a FASTA file of 43 proteins confirmed to exhibit homotypic phase separation behavior that was curated by Vernon *et al*. (27). The UniProt Knowledgebase accession ID (UniProtKB ID), gene name, and primary sequence for the proteins in this set are given in the supporting information (Table S1).

The output includes two sets of plots and four datasets that can be downloaded. The plots are intended to quickly show if the uploaded dataset is enriched or depleted in phase separation potential relative to a reference; here, the reference is the human proteome containing splice variants. Figure 3 reproduces these plots for the Vernon set and shows that this dataset is highly enriched in proteins with long (*N* ≥50) predicted PS IDRs relative to the human proteome (Figure 3A). In addition to length of the predicted PS IDR, the summed classifier distance for every window labeled P has been used as a proxy to estimate the PS potential in a sequence (35). We find that the Vernon set is heavily enriched in proteins with computed PS potentials ≥100, relative to the human proteome (Figure 3B). Also, these data are used to create recall plots from which the area under the curve (AUC) is calculated. AUC values >0.5 in either metric indicate a set of sequences enriched in phase separation potential relative to the reference human proteome.

**Figure 3.**
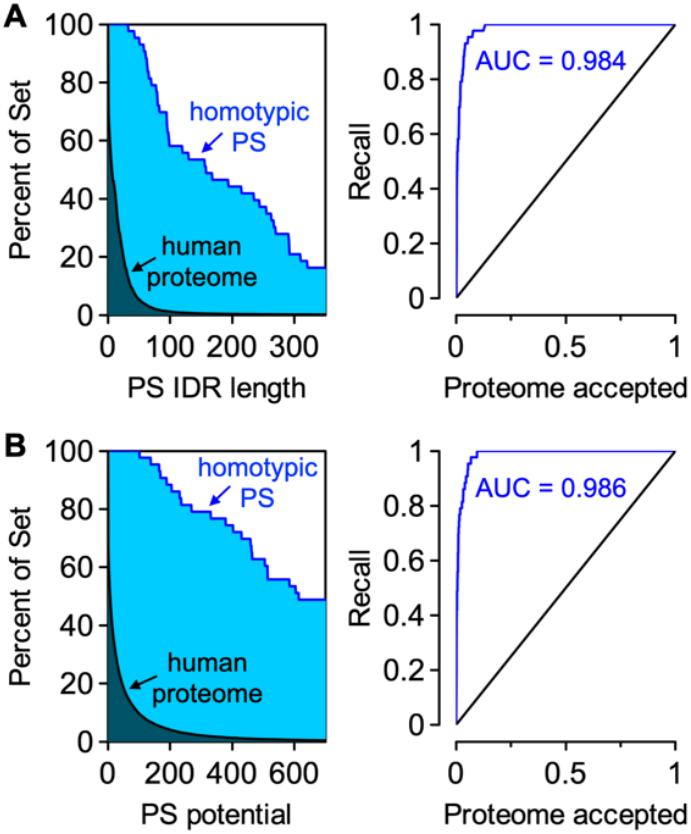
Homotypic PS proteins have long predicted PS IDRs and relatively high computed PS potentials. **A)** ParSe 2.0 finds regions within proteins that are >90% labeled P, which are predicted to be PS IDRs. Left plot: the y-axis is the percent of proteins in a set with PS IDRs at least as long as the length indicated by the x-axis; the human proteome (black) is compared to the Vernon set (blue). **B)** The summed classifier distance of windows labeled P is used to estimate the PS potential in a sequence. Left plot: the y-axis is the percent of proteins in a set with a PS potential equal to or greater than the value indicated by the x-axis. Right plots: in both A and B, the recall plot compares the percent of set for the human proteome against itself (black) and against the Vernon set (blue). AUC is the area under the curve for the Vernon set.

The results can be analyzed using any of four tables, linked after the plots described above. The first is a summary table of sequence-calculated values that can be sorted within the application (or downloaded and sorted separately). Sorting this table by the third column ranks the input file sequences by their computed PS potential (i.e., the classifier distance sum of windows labeled P), or by the fourth column to rank by length of the longest predicted PS IDR (Figure 4). Thus, proteins within the input file predicted to have PS IDRs can be quickly identified by simply sorting the third or fourth columns of this table. This table for the Vernon set is reproduced in the supporting information (Table S2). If the description line for each sequence is formatted according to UniProt, where the line preceding each primary sequence lists the UniProtKB ID followed by the gene name and protein name, these identifying labels will be listed in the last three columns of the summary table.

**Figure 4.**
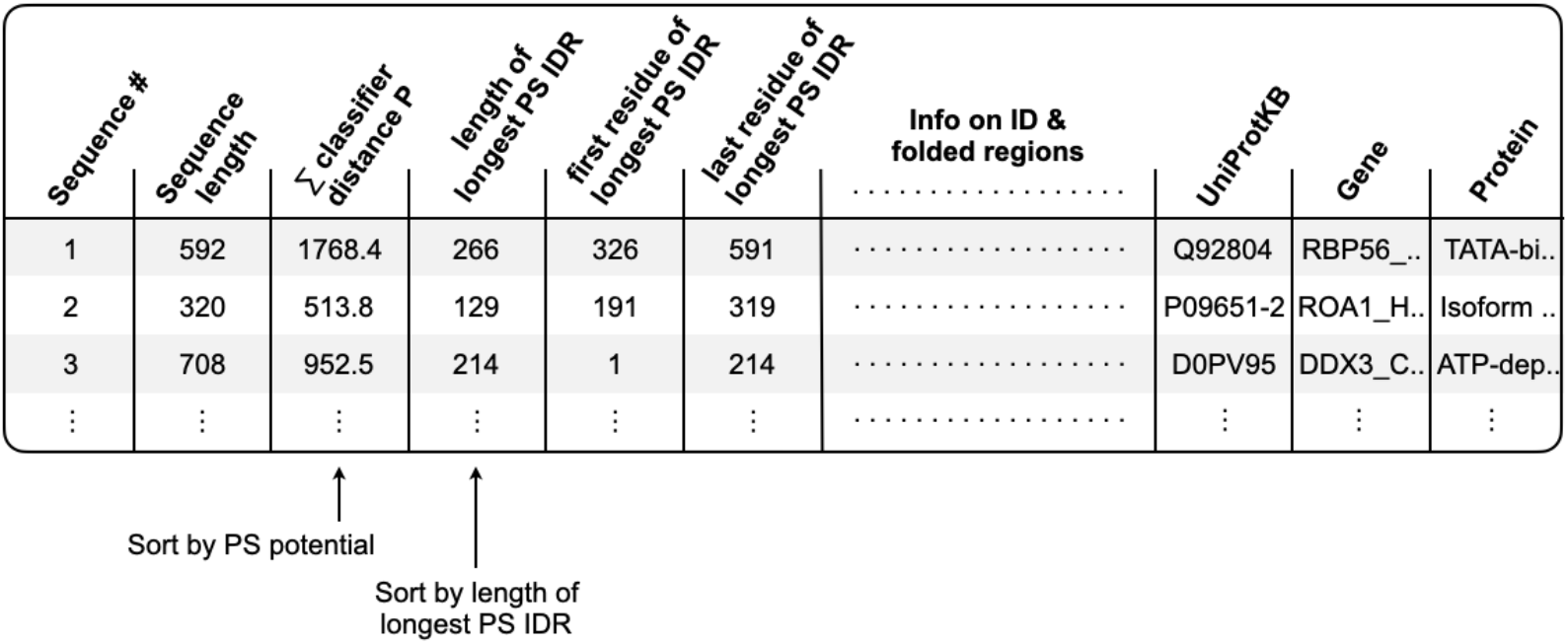
ParSe 2.0 output Summary Table. A key output of the web application is a summary table that lists the longest PS ID, ID, and folded regions predicted to reside within each primary sequence of the user-supplied input file. Sorting this table by the third or fourth columns can be used to quickly identify proteins predicted by the algorithm to exhibit phase separation behavior.

Located below the summary table in the application are tables containing the data used to make the plots described above, allowing for their reproduction outside of the application. Lastly, the application also outputs a FASTA file containing the predicted PS IDRs with length ≥50 that were found within the original input file.

### Comparing ParSe 2.0 to other sequence-based predictors

Previously, we found that ParSe 2.0 is at least as accurate in identifying proteins and regions within proteins that drive phase separation compared to other published phase separation predictors and using publicly available datasets (35). To demonstrate such predictor comparisons here, we used ParSe 2.0 to generate three separate sequence sets derived from the human proteome. Each set contains protein regions of at least 50 residues identified by our algorithm; the sets differ in the distribution of anticipated protein classes. Predicted PS IDRs comprise the first set, predicted IDRs (non-phase-separating) the second set, and predicted folded regions the third.

Of the protein regions ParSe 2.0 predicts to be PS IDRs or IDRs, Figure 5A shows that metapredict (56) and flDPnn (57) also predict those regions as ID for >95% of the sequences found in either set. Metapredict was trained on consensus disorder data from 8 different disorder predictors, whereas flDPnn produced the most accurate predictions of disorder in the recently completed Critical Assessment of protein Intrinsic Disorder (CAID) prediction experiment (58). These data suggest that IDR predictions by ParSe 2.0 are likely to be regions predicted as ID by metapredict and flDPnn. For the set not expected to be ID, only a few (<15%) of the ParSe-predicted folded regions were classified as ID by either of these two ID predictors. Thus, ParSe 2.0, despite predicting IDRs from a single metric, *ϕ*, shows good overall agreement with these two ID predictors when applied to the human proteome.

**Figure 5.**
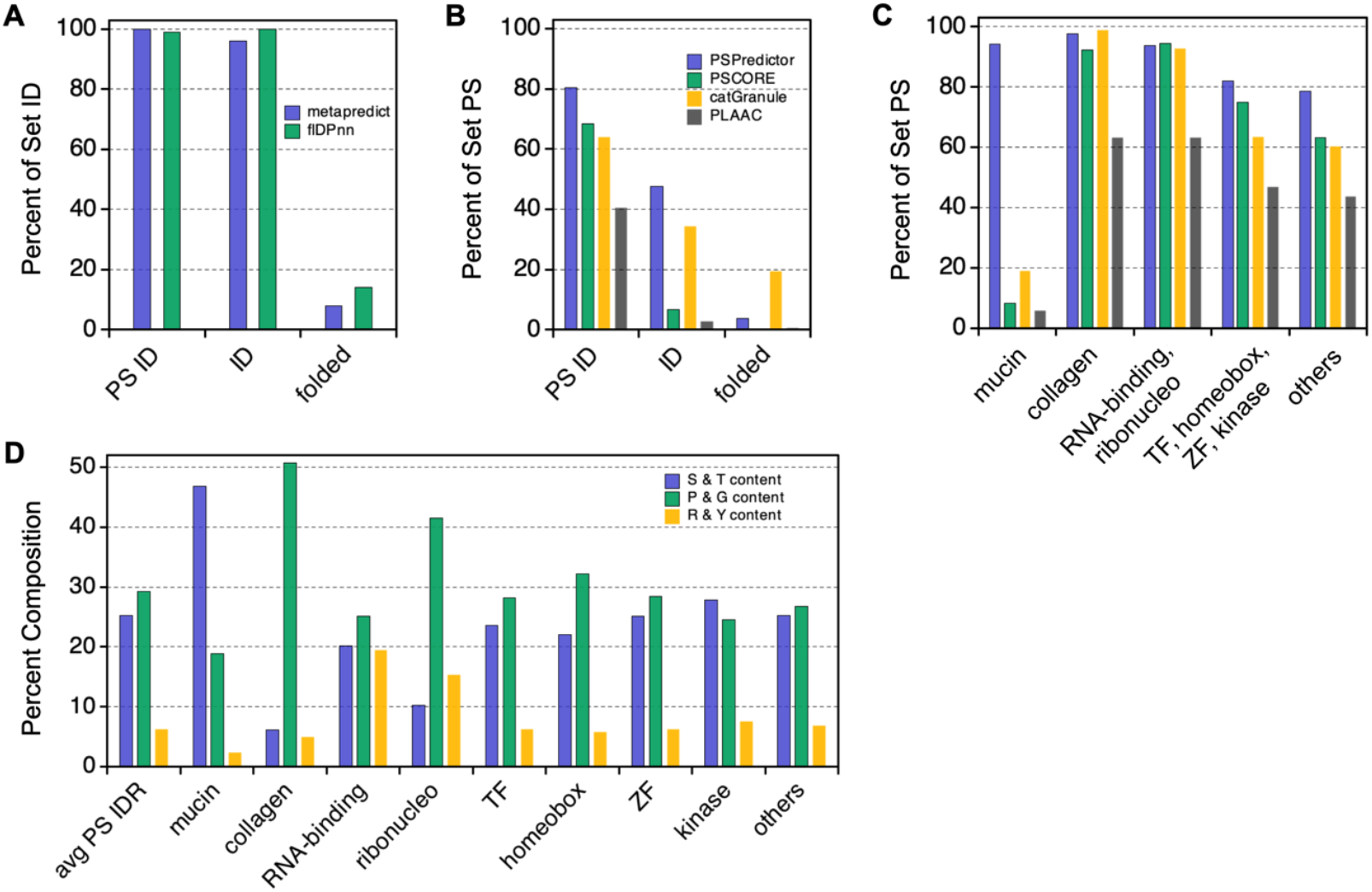
Comparing sequence-based predictors. ParSe 2.0 was used to search the human proteome for protein regions predicted to be PS ID, ID, or folded. Percent of sequences in each set that were **A)** predicted to be ID by metapredict (blue) and flDPnn (green), or **B)** predicted to potentially phase separate by PSPredictor (blue), PScore (green), catGranule (orange), and PLAAC (black). Percent of set values for PSPredictor, catGranule, and flDPnn represent the average from randomly sampling in cumulative 500 primary sequences owing to their relatively low input limits (100, 100, and 20 primary sequences, respectively). **C)** Sequences in the PS IDR set were separated by protein name, creating subsets based on if the name included “mucin”, “collagen”, “RNA-binding”, “ribonucleo”, “transcription factor” (TF), “homeobox”, “zinc finger” (ZF), or “kinase”. RNA-binding and ribonucleo subsets were combined; the TF, homeobox, zinc finger, and kinase subsets also were combined; “others” refers to PS IDR sequences belonging to all other proteins. The percent of each subset predicted to potentially phase separate is shown using the same coloring scheme and predictors as used in panel B. **D)** Overall average percent composition for sequences in the human PS IDR set compared to the subsets for serine and threonine (blue), proline and glycine (green), and arginine and tyrosine (orange).

Figure 5B shows that the predictors PSPredictor (37), PScore (27), catGranule (38), and PLAAC (39) predict phase separation behavior primarily, though not exclusively, in sequences found in the PS IDR set. PSPredictor was developed from machine learning tools that were trained using sequence data of proteins known to phase separate. PScore was developed based on a specific molecular mechanism thought to drive phase separation; the propensity of cation-π and π-π interactions to form cohesive protein interactions. Originally, PLAAC was developed to identify prions, and catGranule targets ID and RNA binding ability; however, these two algorithms have been widely used as proxies for potential phase separation behavior (36, 59, 60). Interestingly, both PSPredictor and catGranule find substantial proportions (>30%) of the predicted non-phase-separating IDR set as possibly showing phase separation behavior.

A key difference among this set of predictors is that ParSe 2.0 and PSPredictor both find PS IDRs within mucins that are mostly missed by PScore, catGranule, and PLAAC (Figure 5C). Mucins are heavily glycosylated proteins that are known to form gel-like assemblies (61). On average, a ParSe-predicted mucin PS IDR is enriched in serine and threonine content and depleted in proline, glycine, arginine, and tyrosine, when its composition is compared to the typical PS IDR predicted within a human protein (Figure 5D). Whether or not ParSe-predicted mucin PS IDRs indeed drive physiological phase separation has not been tested. Interestingly, ParSe-predicted PS IDRs within human RNA-binding and ribonucleoproteins are enriched in arginine and tyrosine content, when compared to the human PS IDR composition average, whereas transcription factors, zinc finger proteins, kinases, and proteins with “homeobox” in their name are found to generally match the average predicted PS IDR composition in humans. Others have found that phase-separating, RNA-binding proteins often have tyrosine-rich, low sequence complexity, prion-like domains and arginine-rich RNA-binding domains (62). ParSe-predicted PS IDRs within collagen proteins are highly enriched in proline and glycine, which is consistent with the atypical amino acid composition of this protein type.

### Modeling mutation effects on phase separation potential

Extensive mutagenesis studies involving several proteins have been used to understand the sequence features that drive phase separation (2, 27, 28, 62–64). The results of those studies implicate specific interactions between amino acids in the formation of phase-separated droplets, e.g., cation-anion, cation-π, and π-π. Overall, the main result of many studies is that multiple, redundant molecular mechanisms contribute to the formation of phase-separated droplets from IDRs (35, 40, 65).

Because the PS potential computed by ParSe 2.0 does not include the effects of pairwise interactions involving combinations of amino acid types, the calculation was expanded to contain both the classifier distance sum of P-labeled positions and terms quantifying the effects of interactions between amino acids, termed *U*_*π*_ for π-π and cation-π interactions and *U*_*q*_ for charge-based effects (35). We trained *U*_*π*_ and *U*_*q*_ against existing data on mutant sequences from Ddx4, LAF1, and A1 (2, 28, 63, 64). However, the different studies used different metrics to quantify phase separation potential. We used the saturation concentration (*c*_*sat*_) at 4 °C and thermodynamic properties associated with phase separation behavior (standard molar Δ*h*°, Δ*s*°, and Δ*g*°) to separately train the calculation. We found that the summed P classifier distance was only moderately able to predict the effects of mutations designed to perturb phase separation behavior. In contrast, the expanded PS potential including *U*_*π*_ and *U*_*q*_ obtained reasonable predictive power, highlighting the importance of pairwise interactions in modulating phase separation behavior (35).

The ParSe 2.0 application targeted at individual protein sequences (accessed at https://stevewhitten.github.io/Parse_v2_web) outputs the computed PS potential both with and without the *U*_*π*_ and *U*_*q*_ extensions. We used this modified algorithm to assess a newly published mutant dataset that was not included in the training of *U*_*π*_ and *U*_*q*_. Mittal and coworkers measured the effects on *c*_*sat*_ at 37 °C from mutation in an artificial IDP consisting of 25-repeats of GRGDSPYS (40). Figure 6 shows that including *U*_*π*_ and *U*_*q*_ in the calculation increased the correlation between experimental *c*_*sat*_ and predicted PS potential (from r = 0.24 to 0.59). If we used *U*_*π*_ and *U*_*q*_ trained previously by Δ*h*°, rather than trained by *c*_*sat*_, the predictive power for capturing sequence effects on *c*_*sat*_ in the new mutant dataset was not as good (r = 0.46, Figure S1). This result is consistent with the observation that mutant rank order in *c*_*sat*_ (at a given temperature) does not necessarily agree with mutant rank order in the measured thermodynamic properties associated with protein phase separation (64).

**Figure 6.**
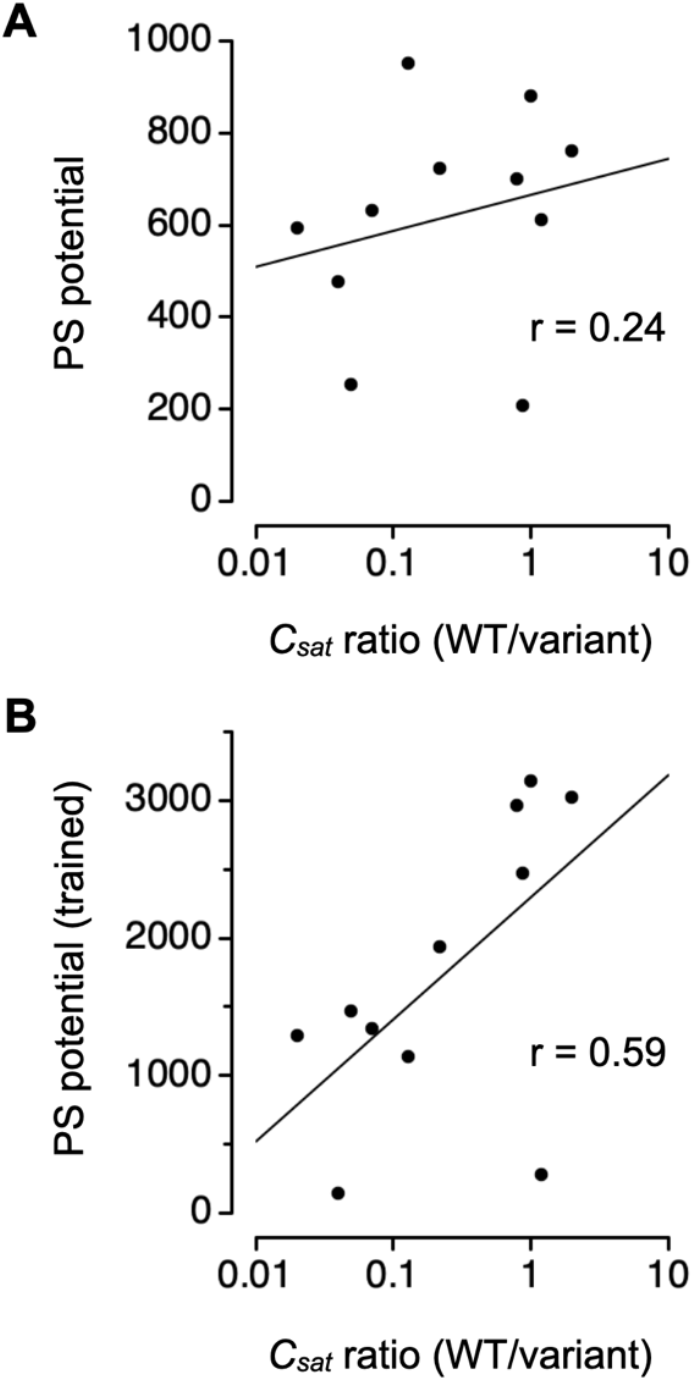
Mutation effects on experimental *c*_*sat*_ compared to the sequence-calculated PS potential. **A)** PS potential calculated as the summed P classifier distance. **B)** Expanded PS potential that includes the summed P classifier distance plus *U*_*π*_ and *U*_*q*_ trained previously using *c*_*sat*_ from a mutant dataset. In both plots, experimental *c*_*sat*_ ratio (x-axis) was digitally extracted from Figure S18A in Rekhi *et al*. (40). WT refers to wild type.

## Discussion

ParSe 2.0 was developed with a particular focus on predicting which IDRs in a protein sequence can lead to phase separation. Our approach for identifying potential PS IDRs is based primarily on sequence composition and not on sequence patterns or combinations of amino acids. This approach was inspired by the finding that a wide variety of amino acid scales show statistically significant differences between curated ID and PS ID datasets, indicating that PS IDRs are a robustly different class of protein region than conventional, non-phase-separating IDRs (35). Similarly, we and others (66–69) have shown that ID (both conventional and PS) and folded protein regions are robustly different in their intrinsic properties, which enables the sequence-based prediction of the modular organization within a protein with respect to ID, PS ID, and folded regions (Figure 1D).

However, to yield reasonable predictive power for mutations that have been designed to test the role of specific amino acid types in driving protein phase separation, the PS potential as computed by ParSe 2.0 has been modified to account for interactions between amino acids. With this modification (i.e., including both *U*_*π*_ and *U*_*q*_), we have been able to match existing data on mutant sequences (35). The original training of *U*_*π*_ and *U*_*q*_ used protein constructs that were based on sequences from Ddx4, LAF1, and A1 (2, 28, 63, 64). The mutant set in Figure 6 is from an artificial protein construct (40) and shows surprisingly good agreement between changes in the computed PS potential and changes in the measured *c*_*sat*_ at 37 °C, especially when the PS potential was modified to include *U*_*π*_ and *U*_*q*_.

We have built web application versions of ParSe 2.0 for the scientific community. Because of its speed, the ParSe 2.0 algorithm can be applied to datasets of large size (Figure 2). The strong performance of ParSe 2.0 on existing datasets, the robust nature of the differences between PS IDRs and conventional IDRs, and the high correlation between ParSe 2.0 and other predictors on databases of PS proteins (35) all give confidence that the algorithm can identify PS IDRs with significant accuracy.

## Methods

### Window calculation of *ϕ*, α, and *v*_*model*_

*ϕ* was calculated as the sequence sum divided by the length, *N*, using the hydrophobicity scale from Bastolla *et al*. (41). For a window of 25 residues, *N* = 25. Similarly, α was calculated as the sequence sum divided by *N* using the α-helix propensity scale from Tanaka and Scheraga (42). *v*_*model*_, introduced previously (30), was calculated by,

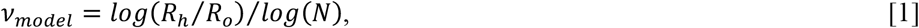

where *R*_*o*_ was a constant set to 2.16 Å, and the hydrodynamic radius, *R*_*h*_, was calculated from sequence using an equation found to be accurate for monomeric IDPs (70–74). The equation to calculate *R*_*h*_ for a disordered sequence is,

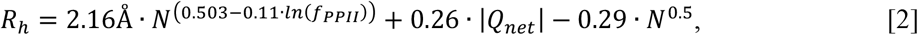

where *f*_*PPII*_ is the fractional number of residues in the PPII conformation, and *Q*_*net*_ is the net charge. *f*_*PPII*_ was estimated from ∑ *P*_*PPII,i*_/*N*, where *P*_*PPII,i*_ is the experimental PPII propensity determined for amino acid type *i* in unfolded peptides by Elam *et al*. (75). *Q*_*net*_ was determined from the number of lysine and arginine residues minus the number of glutamic acid and aspartic acid.

### ParSe 2.0 algorithm

For an arbitrary sequence, whereby the amino acids are restricted to the 20 common types, ParSe 2.0 first reads the sequence to determine its length, *N*. Next, the algorithm uses a sliding window scheme (Fig. 1A) to calculate *v*_*model*_, α-helix propensity, and *ϕ* for every 25-residue segment of the primary sequence. This window scheme can be applied to proteins with *N* >25. A window is labeled F if *ϕ* >0.08 (Fig. 1B). If *ϕ* <0.08, a window is labeled P or D depending on the values of *v*_*model*_ and α-helix propensity. Windows with high α-helix propensity and high *v*_*model*_ are labeled D, while those with low α-helix propensity and low *v*_*model*_ are labeled P. The P/D boundary is given by *v*_*model*_ = -0.244·α-helix propensity + 0.789. The window label is assigned to the central residue in that window. N- and C-terminal residues not belonging to a central window position are assigned the label of the central residue in the first and last window, respectively, of the whole sequence. Protein regions predicted to be PS, ID, or folded are determined by finding contiguous residue positions of length ≥20 that are ≥90% of only one label P, D, or F, respectively.

### Classifier distance calculation

The classifier distance is the normalized distance of a ParSe 2.0 generated window into its classifier sector (i.e., F, D, or P sector) and relative to the cutoff boundary (Fig. 1B). For F-labeled windows, the classifier distance is *ϕ* (of the window) minus the cutoff value of 0.08 and then normalized to distance of the folded training set mean *ϕ* (0.1164) to the cutoff. Specifically, this is (*ϕ* – 0.08)/ (0.1164–0.08). For P or D labeled windows, first we find the point on the P/D boundary (defined above) that makes a perpendicular bisector when paired with the window values of *v*_*model*_ and α-helix propensity. Then the distance between this point and the point defined by the window values of *v*_*model*_ and α-helix propensity is determined. Specifically, this distance is sqrt((α – x)·(α – x) + (*v*_*model*_ – y)·(*v*_*model*_ – y)), where α and *v*_*model*_ are defined above, x is (α/0.244 + 0.789 – *v*_*model*_)/(0.244 + 1/0.244), and y is (x – α)/0.244 + *v*_*model*_. This distance is normalized by dividing by 0.019, the distance from the boundary to either of the training set means.

### Computed PS potential

The PS potential for a sequence is the summed classifier distance for every window labeled P. This potential can be expanded to include contributions of aromatic and cation-π interactions (*U*_*π*_) and charge-based interactions (*U*_*q*_) as described below.

### Contribution of aromatic and cation-π interactions to the PS potential

The contributions of aromatic and cation-π interactions to protein phase separation follows the observed rank order by Wang *et al*.: Tyr-Arg > Tyr-Lys ∼ Phe-Arg > Phe-Lys (62). To mimic this ranking, we assumed 3:2:1 weighting and, also, that Phe–Tyr interactions would contribute comparably to Phe–Lys interactions,

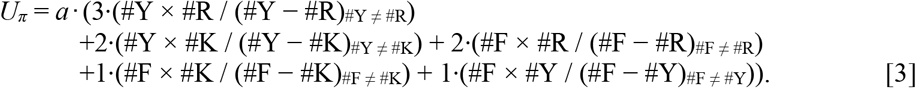

In equation 3, #Y, #R, #F, and #K represent the number of Tyr, Arg, Phe, and Lys residues, respectively, calculated on a per-window basis. Thus, *U*_*π*_ increases with increasing Tyr, Arg, Phe, and Lys content and more so when interaction partners are present at similar levels. When the divisor is zero (e.g., when #Y = #R), it is changed to 1 to avoid infinite potentials. The fitting parameter *a* was determined previously (35) by finding the optimal correlation of the expanded PS potential to experimental Δ*h°* (finding *a* = 0.14), Δ*s°* (finding *a* = 0.08), Δ*g°* (finding *a* = 0.11), or *c*_*sat*_ at 4 °C (finding *a* = 0.28). Window-specific *U*_*π*_ is added to the classifier distance at windows labeled P. *U*_*π*_ also is calculated at D-labeled windows, allowing for the possibility of labels changing from D to P. This would occur when the value for *U*_*π*_ was larger than the classifier distance at a D-labeled window. Thus, protein regions that otherwise have characteristics more like the ID set, in *v*_*model*_ and α-helix propensity, could be labeled P if *U*_*π*_ was large enough. Here, the given classifier distance was determined by the difference between *U*_*π*_ and the original classifier distance of the window formerly labeled D.

### Contribution of charge-based interactions to the PS potential

The contributions of charge-based interactions to protein phase separation follows the observations by Schuster *et al*. (63) and Bremer *et al*. (64) that changes in the sequence charge decoration, *SCD*, and net charge per residue, *NCPR*, respectively, can affect phase separation potential. Thus, a simple charge-based potential was defined,

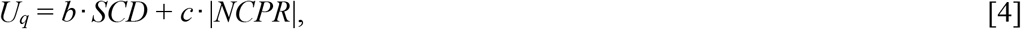

where *b* and *c* are fitting parameters, and *U*_*q*_ is calculated on a per-window basis. *U*_*q*_ is added to the classifier distance at each window labeled P and is applied to windows labeled D, following the scheme described above for *U*_*π*_, again allowing for the possibility of labels changing from D to P. The parameters *b* and *c* were determined previously (35) by finding the optimal correlation of the expanded PS potential and Δ*h°* (finding 8.4 and 5.6, respectively), Δ*s°* (finding 4.6 and 7.0, respectively), Δ*g°* (finding 5.2 and 5.4, respectively), or *c*_*sat*_ at 4 °C (finding -16.0 and 33, respectively). *NCPR* is the number of Lys and Arg residues minus the number of Glu and Asp residues, divided by *N. SCD* is calculated by *N*^-1^∑_*i*_∑_*j,j>i*_(*q*_*i*_*q*_*j*_)|*j-i*|^1/2^, where *q* is the amino acid– specific charge (76).

### Metapredict calculation

Metapredict score (56), which predicts the presence of ID in a sequence, was calculated by computer algorithm using the Python script available at http://metapredict.net. The per-residue average metapredict score, when >0.5, was used to classify a protein region as predicted to be ID.

### flDPnn calculation

flDPnn score (57), which predicts the presence of ID in a sequence, was calculated by using the webserver available at http://biomine.cs.vcu.edu/servers/flDPnn. The per-residue average flDPnn binary score, when >0.5, was used to classify a protein region as predicted to be ID.

### PSPredictor calculation

PSPredictor score was calculated by using the PSPredictor (37) webtool available at http://www.pkumdl.cn:8000/PSPredictor. PSPredictor score, when >0.5, was used to classify a protein region as predicted to exhibit phase separation behavior.

### PSCORE calculation

PSCORE, which is a phase separation propensity predictor (27), was calculated by computer algorithm using the Python script and associated database files available at https://doi.org/10.7554/eLife.31486.022. The overall PScore, when >4, was used to classify a protein region as predicted to exhibit phase separation behavior.

### Granule propensity calculation

Granule propensity was calculated by using the catGranule (38) webtool available at http://www.tartaglialab.com. Granule propensity, when >0, was used to classify a protein region as predicted to exhibit phase separation behavior.

### PLAAC LLR calculation

LLR score, which identifies prion-containing sequences (39), was calculated by using the webtool available at http://plaac.wi.mit.edu. The LLR score, when >0, was used to classify a protein region as predicted to exhibit phase separation behavior.

## Data availability

A web application of ParSe 2.0 that evaluates individual protein sequences can be accessed at https://stevewhitten.github.io/Parse_v2_web. A web application of ParSe 2.0 that can be used to quickly find phase-separating proteins within large sequence sets can be accessed at https://stevewhitten.github.io/Parse_v2_FASTA. The source code for both applications can be accessed at https://github.com/stevewhitten.

## Author contributions

L. E. H. and S. T. W. conceptualization; C. W. and S. T. W., programming; K. A. L., N. C. F., L. E. H., and S. T. W., methodology; K. A. L., N. C. F., L. E. H., and S. T. W., formal analysis; S. T. W., writing–original draft; K.A.L., N. C. F., L. E. H, and S. T. W., writing– review and editing.

## Funding and additional information

This work was supported by the National Institutes of Health under grants R35GM119755 (L. E. H.), and R01AI139479 (N. C. F.), the National Science Foundation under grants 1818090 (N. C. F.) and 1943488 (L. E. H.), and Texas State University Office of Research and Sponsored Projects through the Research Enhancement Program (S. T. W. and K. A. L.). No nongovernmental sources were used to fund this project. The content is solely the responsibility of the authors and does not necessarily represent the official views of the NSF or NIH.

