## Supplemental Information for "ParSe 2.0: A web application that enables proteome-scale searches for sequences that drive protein-mediated phase separation"

### Contents:

Supporting Tables
S1. Proteins confirmed to exhibit homotypic phase separation behavior.
S2. Summary table for the set of proteins that exhibit homotypic phase separation behavior.

Supporting Figures
S1. Mutation effects on experimental $c_{sat}$ compared to the PS potential trained previously using experimental $\Delta h°$ from a mutant dataset.

Supporting References

### Supporting Tables

#### Table S1. Proteins confirmed to exhibit homotypic phase separation behavior.

| UniProtKB ID [a] | Gene name | Primary sequence |
|---|---|---|
| Q92804 | RBP56_HUMAN | MSDSGSYGQSGGEQQSYSTYGNPGSQGYGQASQSYSGYGQTTDSSYGQ<br>NYSGYSSYGQSQSGYSQSYGGYENQKQSSYSQQPYNNQGQQQNMESSG<br>SQGGRAPSYDQPDYGQQDSYDQQSGYDQHQGSYDEQSNYDQQHDSYSQ<br>NQQSYHSQRENYSHHTQDDRRDVSRYGEDNRGYGGSQGGGRGRGGYDK<br>DGRGPMTGSSGGDRGGFKNFGGHRDYGPRTDADSESDNSDNNTIFVQG<br>LGEGVSTDQVGEFFKQIGIIKTNKKTGKPMINLYTDKDTGKPKGEATV<br>SFDDPPSAKAAIDWFDGKEFHGNIIKVSFATRRPEFMRGGGSGGGRRG<br>RGGYRGRGGFQGRGGDPKSGDWVCPNPSCGNMNFARRNSCNQCNEPRP<br>EDSRPSGGDFRGRGYGGERGYRGRGGRGGDRGGYGGDRSGGGYGGDRS<br>SGGGYSGDRSGGGYGGDRSGGGYGGDRGGGYGGDRGGGYGGDRGGGYG<br>GDRGGYGGDRGGGYGGDRGGYGGDRGGYGGDRGGYGGDRGGYGGDRSR<br>GGYGGDRGGGSGYGGDRSGGYGGDRSGGGYGGDRGGGYGGDRGGYGGK<br>MGGRNDYRNDQRNRPY |
| P09651-2 | ROA1_HUMAN | MSKSESPKEPEQLRKLFIGGLSFETTDESLRSHFEQWGTLTDCVVMRD<br>PNTKRSRGFGFVTYATVEEVDAAMNARPHKVDGRVVEPKRAVSREDSQ<br>RPGAHLTVKKIFVGGIKEDTEEHHLRDYFEQYGKIEVIEIMTDRGSGK<br>KRGFAFVTFDDHDSVDKIVIQKYHTVNGHNCEVRKALSKQEMASASSS<br>QRGRSGSGNFGGGRGGGFGGNDNFGRGGNFSGRGGFGGSRGGGGYGGS<br>GDGYNGFGNDGSNFGGGGSYNDFGNYNNQSSNFGPMKGGNFGGRSSGP<br>YGGGGQYFAKPRNQGGYGGSSSSSSYGSGRRF |
| D0PV95 | DDX3_CAEEL | MESNQSNNGGSGNAALNRGGRYVPPHLRGGDGGAAAAASAGGDDRRGG<br>AGGGGYRRGGGNSGGGGGGGYDRGYNDNRDDRDNRGGSGGYGRDRNYE<br>DRGYNGGGGGGGNRGYNNNRGGGGGGYNRQDRGDGGSSNFSRGGYNNR<br>DEGSDNRGSGRSYNNDRRDNGGDGQNTRWNNLDAPPSRGTSKWENRGA<br>RDERIEQELFSGQLSGINFDKYEEIPVEATGDDVPQPISLFSDLSLHE<br>WIEENIKTAGYDRPTPVQKYSIPALQGGRDLMSCAQTGSGKTAAFLVP<br>LVNAILQDGPDAVHRSVTSSGGRKKQYPSALVLSPTRELSLQIFNESR<br>KFAYRTPITSALLYGGRENYKDQIHKLRLGCHILIATPGRLIDVMDQG<br>LIGMEGCRYLVLDEADRMLDMGFEPQIRQIVECNRMPSKEERITAMFS<br>ATFPKEIQLLAQDFLKENYVFLAVGRVGSTSENIMQKIVWVEEDEKRS<br>YLMDLLDATGDSSLTLVFVETKRGASDLAYYLNRQNYEVVTIHGDLKQ<br>FEREKHLDLFRTGTAPILVATAVAARGLDIPNVKHVINYDLPSDVDEY<br>VHRIGRTGRVGNVGLATSFFNDKNRNIARELMDLIVEANQELPDWLEG<br>MSGDMRSGGGYRGRGGRGNGQRFGGRDHRYQGGSGNGGGGNGGGGGFG<br>GGGQRSGGGGGFQSGGGGGRQQQQQQRAQPQQDWWS |
| H3BNZ4 | H3BNZ4_HUMAN | MASNDYTQQATQSYGAYPTQPGQGYSQQSSQPYGQQSYSGYSQSTDTS<br>GYGQSSYSSYGQSQNTGYGTQSTPQGYGSTGGYGSSQSSQSSYGQQSS<br>YPGYGQQPAPSSTSGSYGSSSQSSSYGQPQSGSYSQQPSYGGQQQSYG<br>QQQSYNPPQGYGQQNQYNSSSGGGGGGGGGGNYGQDQSSMSSGGGSGG<br>GYGNQDQSGGGGSGGYGQQDRGGRGRGGSGGGGGGGGGGYNRSSGGYE<br>PRGRGGGRGGRGGMGPSGPRITS |
| O00571 | DDX3X_HUMAN | MSHVAVENALGLDQQFAGLDLNSSDNQSGGSTASKGRYIPPHLRNREA<br>TKGFYDKDSSGWSSSKDKDAYSSFGSRSDSRGKSSFFSDRGSGSRGRF<br>DDRGRSDYDGIGSRGDRSGFGKFERGGNSRWCDKSDEDDWSKPLPPSE<br>RLEQELFSGGNTGINFEKYDDIPVEATGNNCPPHIESFSDVEMGEIIM<br>GNIELTRYTRPTPVQKHAIPIIKEKRDLMACAQTGSGKTAAFLLPILS<br>QIYSDGPGEALRAMKENGRYGRRKQYPISLVLAPTRELAVQIYEEARK<br>FSYRSRVRPCVVYGGADIGQQIRDLERGCHLLVATPGRLVDMMERGKI<br>GLDFCKYLVLDEADRMLDMGFEPQIRRIVEQDTMPPKGVRHTMMFSAT<br>FPKEIQMLARDFLDEYIFLAVGRVGSTSENITQKVVWVEESDKRSFLL<br>DLLNATGKDSLTLVFVETKKGADSLEDFLYHEGYACTSIHGDRSQRDR<br>EEALHQFRSGKSPILVATAVAARGLDISNVKHVINFDLPSDIEEYVHR<br>IGRTGRVGNLGLATSFFNERNINITKDLLDLLVEAKQEVPSWLENMAY<br>EHHYKGSSRGRSKSSRFSGGFGARDYRQSSGASSSSFSSSRASSSRSG<br>GGGHGSSRGFGGGGYGGFYNSDGYGGNYNSQGVDWWGN |
| Q9NQI0 | DDX4_HUMAN | MGDEDWEAEINPHMSSYVPIFEKDRYSGENGDNFNRTPASSSEMDDGP<br>SRRDHFMKSGFASGRNFGNRDAGECNKRDNTSTMGGFGVGKSFGNRGF |

| | | |
|---|---|---|
| | | SNSRFEDGDSSGFWRESSNDCEDNPTRNRGFSKRGGYRDGNNSEASGP<br>YRRGGRGSFRGCRGGFGLGSPNNDLDPDECMQRTGGLFGSRRPVLSGT<br>GNGDTSQSRSGSGSERGGYKGLNEEVITGSGKNSWKSEAEGGESSDTQ<br>GPKVTYIPPPPPEDEDSIFAHYQTGINFDKYDTILVEVSGHDAPPAIL<br>TFEEANLCQTLNNNIAKAGYTKLTPVQKYSIPIILAGRDLMACAQTGS<br>GKTAAFLLPILAHMMHDGITASRFKELQEPECIIVAPTRELVNQIYLE<br>ARKFSFGTCVRAVVIYGGTQLGHSIRQIVQGCNILCATPGRLMDIIGK<br>EKIGLKQIKYLVLDEADRMLDMGFGPEMKKLISCPGMPSKEQRQTLMF<br>SATFPEEIQRLAAEFLKSNYLFVAVGQVGGACRDVQQTVLQVGQFSKR<br>EKLVEILRNIGDERTMVFVETKKKADFIATFLCQEKISTTSIHGDREQ<br>REREQALGDFRFGKCPVLVATSVAARGLDIENVQHVINFDLPSTIDEY<br>VHRIGRTGRCGNTGRAISFFDLESDNHLAQPLVKVLTDAQQDVPAWLE<br>EIAFSTYIPGFSGSTRGNVFASVDTRKGKSTLNTAGFSSSQAPNPVDD<br>ESWD |
| Q15056 | IF4H_HUMAN | MADFDTYDDRAYSSFGGGRGSRGSAGGHGSRSQKELPTEPPYTAYVGN<br>LPFNTVQGDIDAIFKDLSIRSVRLVRDKDTDKFKGFCYVEFDEVDSLK<br>EALTYDGALLGDRSLRVDIAEGRKQDKGGFGFRKGGPDDRGMGSSRES<br>RGGWDSRDDFNSGFRDDFLGGRGGSRPGDRRTGPPMGSRFRDGPPLRG<br>SNMDFREPTEEERAQRPRLQLKPRTVATPLNQVANPNSAIFGGARPRE<br>EVVQKEQE |
| P14907 | NSP1_YEAST | MNFNTPQQNKTPFSFGTANNNSNTTNQNSSTGAGAFGTGQSTFGFNNS<br>APNNTNNANSSITPAFGSNNTGNTAFGNSNPTSNVFGSNNSTTNTFGS<br>NSAGTSLFGSSSAQQTKSNGTAGGNTFGSSSLFNNSTNSNTTKPAFGG<br>LNFGGGNNTTPSSTGNANTSNNLFGATANANKPAFSFGATTNDDKKTE<br>PDKPAFSFNSSVGNKTDAQAPTTGFSFGSQLGGNKTVNEAAKPSLSFG<br>SGSAGANPAGASQPEPTTNEPAKPALSFGTATSDNKTTNTTPSFSFGA<br>KSDENKAGATSKPAFSFGAKPEEKKDDNSSKPAFSFGAKSNEDKQDGT<br>AKPAFSFGAKPAEKNNNETSKPAFSFGAKSDEKKDGDASKPAFSFGAK<br>PDENKASATSKPAFSFGAKPEEKKDDNSSKPAFSFGAKSNEDKQDGTA<br>KPAFSFGAKPAEKNNNETSKPAFSFGAKSDEKKDGDASKPAFSFGAKS<br>DEKKDSDSSKPAFSFGTKSNEKKDSGSSKPAFSFGAKPDEKKNDEVSK<br>PAFSFGAKANEKKESDESKSAFSFGSKPTGKEEGDGAKAAISFGAKPE<br>EQKSSDTSKPAFTFGAQKDNEKKTEESSTGKSTADVKSSDSLKLNSKP<br>VELKPVSLDNKTLDDLVTKWTNQLTESASHFEQYTKKINSWDQVLVKG<br>GEQISQLYSDAVMAEHSQNKIDQSLQYIERQQDELENFLDNFETKTEA<br>LLSDVVSTSSGAAANNNDQKRQQAYKTAQTLDENLNSLSSNLSSLIVE<br>INNVSNTFNKTTNIDINNEDENIQLIKILNSHFDALRSLDDNSTSLEK<br>QINSIKK |
| F8WC90 | F8WC90_HUMAN | ASTDYSTYSQAAAQQGYSAYTAQPTQGYAQTTQQAYGQQSYGTYGQPT<br>DVSYTQAQTTATYGQTAYATSYGQPPTGYTTPTAPQAYSQPVQGYGTG<br>AYDTTTATVTTTQASYAAQSAYGTQPAYPAYGQQPAATAPTRPQDGNK<br>PTETSQPQSSTGGYNQPSLGYGQSNYSYPQVPGSYPMQPVTAPPSYPP<br>TSYSSTQPTSYDQSSYSQQNTYGQPSSYGQQSSYGQQSSYGQQPPTSY<br>PPQTGSYSQAPSQYSQQSSSYGQQSSFRQDHPSSMGVYGQESGGFSGP<br>GEN |
| P31483 | TIA1_HUMAN | MEDEMPKTLYVGNLSRDVTEALILQLFSQIGPCKNCKMIMDTAGNDPY<br>CFVEFHEHRHAAAALAAMNGRKIMGKEVKVNWATTPSSQKKDTSSSTV<br>VSTQRSQDHFHVFVGDLSPEITTEDIKAAFAPFGRISDARVVKDMATG<br>KSKGYGFVSFFNKWDAENAIQQMGGQWLGGRQIRTNWATRKPPAPKST<br>YESNTKQLSYDEVVNQSSPSNCTVYCGGVTSGLTEQLMRQTFSPFGQI<br>MEIRVFPDKGYSFVRFNSHESAAHAIVSVNGTTIEGHVVKCYWGKETL<br>DMINPVQQQNQIGYPQPYGQWGQWYGNAQQIGQYMPNGWQVPAYGMYG<br>QAWNQQGFNQTQSSAPWMGPNYGVQPPQGQNGSMLPNQPSGYRVAGYE<br>TQ |
| P15502 | ELN_HUMAN | MAGLTAAAPRPGVLLLLLSILHPSRPGGVPGAIPGGVPGGVFYPGAGL<br>GALGGGALGPGGKPLKPVPGGLAGAGLGAGLGAFPAVTFPGALVPGGV<br>ADAAAAYKAAKAGAGLGGVPGVGGLGVSAGAVVPQPGAGVKPGKVPGV<br>GLPGVYPGGVLPGARFPGVGVLPGVPTGAGVKPKAPGVGGAFAGIPGV<br>GPFGGPQPGVPLGYPIKAPKLPGGYGLPYTTGKLPYGYGPGGVAGAAG<br>KAGYPTGTGVGPQAAAAAAAKAAAKFGAGAAGVLPGVGGAGVPGVPGA<br>IPGIGGIAGVGTPAAAAAAAAAAKAAKYGAAAGLVPGGPGFGPGVVGV<br>PGAGVPGVGVPGAGIPVVPGAGIPGAAVPGVVSPEAAAKAAAKAAKYG<br>ARPGVGVGGIPTYGVGAGGFPGFGVGVGGIPGVAGVPGVGGVPGVGGV<br>PGVGISPEAQAAAAAKAAKYGAAGAGVLGGLVPGAPGAVPGVPGTGGV |

| | | |
|---|---|---|
| | | PGVGTPAAAAAKAAAKAAQFGLVPGVGVAPGVGVAPGVGVAPGVGLAP<br>GVGVAPGVGVAPGVGVAPGIGPGGVAAAAKSAAKVAAKAQLRAAAGLG<br>AGIPGLGVGVGVPGLGVGAGVPGLGVGAGVPGFGAGADEGVRRSLSPE<br>LREGDPSSSQHLPSTPSSPRVPGALAAAKAAKYGAAVPGVLGGLGALG<br>GVGIPGGVVGAGPAAAAAAAKAAAKAAQFGLVGAAGLGGLGVGGLGVP<br>GVGGLGGIPPAAAAKAAKYGAAGLGGVLGGAGQFPLGGVAARPGFGLS<br>PIFPGGACLGKACGRKRK |
| P22626-2 | ROA2_HUMAN | MEREKEQFRKLFIGGLSFETTEESLRNYYEQWGKLTDCVVMRDPASKR<br>SRGFGFVTFSSMAEVDAAMAARPHSIDGRVVEPKRAVAREESGKPGAH<br>VTVKKLFVGGIKEDTEEHHLRDYFEEYGKIDTIEIITDRQSGKKRGFG<br>FVTFDDHDPVDKIVLQKYHTINGHNAEVRKALSRQEMQEVQSSRSGRG<br>GNFGFGDSRGGGGNFGPGPGSNFRGGSDGYGSGRGFGDGYNGYGGGPG<br>GGNFGGSPGYGGGRGGYGGGGPGYGNQGGGYGGGYDNYGGGNYGSGNY<br>NDFGNYNQQPSNYGPMKSGNFGGSRNMGGPYGGGNYGPGGSGGSGGYG<br>GRSRY |
| P22232 | FBRL_XENLA | MRPGFSPRGGRGGFGDRGGFGGRGGFGDRGGFRGGSRGGFGGRGRGGD<br>RGGRGGFRGGFSSPGRGGPRGGGRGGFGGGRGGFGAGRKVIVEPHRHE<br>GIFICRGKEDALVTKNLVPGESVYGEKRISVEDGEVKTEYRAWNPFRS<br>KIAAAILGGVDQIHIKPGVKVLYLGAASGTTVSHVSDVVGPEGLVYAV<br>EFSHRSGRDLINVAKKRTNIIPVIEDARHPHKYRILVGMVDVVFADVA<br>QPDQTRIVALNAHNFLKNGGHFVISIKANCIDSTAAPEAVFAAEVKKM<br>QQENMKPQEQLTLEPYERDHAVVVGIYRPPPKQKK |
| G5EBV6 | PGL3_CAEEL | MEANKRQIVEVDGIKSYFFPHLAHYLASNDELLVNNIAQANKLAAFVL<br>GATDKRPSNEEIAEMILPNDSSAYVLAAGMDVCLILGDDFRPKFDSGA<br>EKLSQLGQAHDLAPIIDDEKKISMLARKTKLKKSNDAKILQVLLKVLG<br>AEEAEEKFVELSELSSALDLDFDVYVLAKLLGFASEELQEEIEIIRDN<br>VTDAFEACKPLLKKLMIEGPKIDSVDPFTQLLLTPQEESIEKAVSHIV<br>ARFEEASAVEDDESLVLKSQLGYQLIFLVVRSLADGKRDASRTIQSLM<br>PSSVRAEVFPGLQRSVFKSAVFLASHIIQVFLGSMKSFEDWAFVGLAE<br>DLESTWRRRAIAELLKKFRISVLEQCFSQPIPLLPQSELNNETVIENV<br>NNALQFALWITEFYGSESEKKSLNQLQFLSPKSKNLLVDSFKKFAQGL<br>DSKDHVNRIIESLEKSSSSEPSATAKQTTTSNGPTTVSTAAQVVTVEK<br>MPFSRQTIPCEGTDLANVLNSAKIIGESVTVAAHDVIPEKLNAEKNDN<br>TPSTASPVQFSSDGWDSPTKSVALPPKISTLEEEQEEDTTITKVSPQP<br>QERTGTAWGSGDATPVPLATPVNEYKVSGFGAAPVASGFGQFASSNGT<br>SGRGSYGGGRGGDRGGRGAYGGDRGRGGSGDGSRGYRGGDRGGRGSYG<br>EGSRGYQGGRAGFFGGSRGGS |
| Q14011 | CIRBP_HUMAN | MASDEGKLFVGGLSFDTNEQSLEQVFSKYGQISEVVVVKDRETQRSRG<br>FGFVTFENIDDAKDAMMAMNGKSVDGRQIRVDQAGKSSDNRSRGYRGG<br>SAGGRGFFRGGRGRGRGFSRGGGDRGYGGNRFESRSGGYGGSRDYYSS<br>RSQSGGYSDRSSGGSYRDSYDSYATHNE |
| D3KYQ3 | D3KYQ3_TETTH | MFGNTGGGGLFGNTQTQQTGGGLFGQPQQTQFGQTGATGGGLFGGATN<br>TFGGGGGGGLFGGNNNQQTNPTAGGGIFGQGTTGLGGAPAQTGGGLFG<br>APQNNQQGGGLFGGGTTTGGGMFGNQANTQTGGGGLFGGPSQPTTQPP<br>AFSLNNPTTGGGGLFGQPANTMGGNNGGLFGGQTNSFGANNNMLGNNN<br>RPQGAGIFGGATTTAPTGNTGMFGGIGANNGGGGLFGMNNTNTNPTGG<br>FGATNPTAGGGGLFGGGATTTGGGGLFGGGNTQGGGLLGTANTTAGGL<br>LGGGFNMNNNTGGILGQTNNQFGLGSFGTNNNAAAAPFQPKASANGVL<br>TKPNEKNLCYAISNGTDFCIFELALTQRKLVKAGQLKPGAQQAGGMFG<br>QPAQGGNGLFGGGGAATTTPFGGAQNGNLFGGQNTQAQGGGLFGAPVN<br>NAATGAGGGLFGAKPAATTTGGGLFGQMPAQTGGFLGNTATQPAGGGL<br>FGGATTTQAPGGGGGGGLFGGNTTAATTGGGLFGGNTQTGGATGGLFG<br>GQQPNNQGGLFLNTGNANNANTGGGLFGGATTTPATGGGLFGGSTNTQ<br>PGLATGGGLFGNNQGASQPAAQGGLFGGAAPQQNSLFGGATAGGQTGG<br>LFGGATGATQQQGGGLFGQTASNPTQGGGLFGAANPGLGGAAAGTQVA<br>QPGLNGFQSLIQLLEQTQNQNNDKFALDDDTEQNWVYGKLRDGYQGGS<br>LGYFQGGDSYSQSGNKFIQLEQQSLYGYQKKKQAYNQFKNLAHRDEKE<br>KKDEMKKLLSSFSSSFVGIGSEQKKQVNLSSQSQALPTRLQQRKLSGS<br>SEISHVSEKLKQIEKSDLCIDFIKSRTSNRIRIPVKPRYTFGDVKQKI<br>LNELINDNEINVRNFRLQARLKGPNKAPKPLRDTDPVFQNKNLPFENI<br>DYIELVERSGNISSLNATLNNNNNSFVLNNIPNDEIDDRNSLQNNSRP<br>IDQKPLLKARSQRINLPFRFSSMPSVDEMEKFTEQQLSQVENFVLENE<br>YARIVFQEPVNLYNLDLSQLIFEESRISLEQVPESDRLNKKCIIEFKK |

| | | |
|---|---|---|
| | | FGLKQLNSGKSKEEVKTQVERLVKKKNLQLVKPYDVDNTKLVYIQDKF<br>Y |
| Q387F2 | Q387F2_TRYB2 | MSAGFGGGFGQPAATGFGQQPTGGFGQAPQGGAFGQVAPAATGFGQPS<br>QSAVTGGFGQTNTGGFGQPAATGFGQPAQGAVTGGFGQTNTGGFGQPA<br>ATGFGQPAQSAVTGGFGQTNTGGFGQPAQGGFGQTAAAANAFGQAGPS<br>GGFGQTNTGGFGQQSNSGFGQAGRGATAGFGQPGTGFGQPATGGFGQA<br>TSASPFGQAAAGRGVGGGFGTAAGTVGGFGQPAATGGFGQTATTGGFG<br>QPAQGAAAGGFGQPATGGFGQATSASPFGQAAAGRGVGGGFGTAAGTV<br>GGFGQPAAPGGFGQTATAGGFGQPARGAAAGGFGQPATGGFGQATSAS<br>PFGQAAAGRGVGGGFGTAAGTVGGFGQPAATGGFGQTATTGGFGQPAQ<br>GANTFGQGTPSAGGFGQAGRGVTGGFGQTGVTGGFGQTATTGGFGQPA<br>QGAATGGFGQAGRGAADGFGRPAQGAAAGGFGQPATGGFGQATSASPF<br>GQAAAGRGVGGGFGTAAGTVGGFGQPAAPGGFGQTATTGGFGQPGRGA<br>AAGGFGQPATGGLGLAGGSGFGAAAGAGGFGQQSTASPWGGGATPAAP<br>VDVSLLNLPEFADKPYGNVLLFAPEKPPKKPISVGHTPSPAAPFNPIS<br>VPRYQQRIPVGIPAPTVSANVKLQPTPFSTSALSSVELRELLNASKVN<br>LGVSDDVQKIDSPKVSRVVAAGEKPPSHDAAVFAPVCSKEEYILDPPL<br>ATLQGLAVRQLQEVYGFSVYRRDGKCSVHFLEPVNLVRCDIAEIVELR<br>PSGEVKLYPCVQNPPSIGQGLNVKARVTVNGVIGTTSNDLLIRCQKEG<br>NRFESYDVNTGTWVYVMNVGDDGQNEYDDADREVDIVETDTQVQDGEQ<br>FPVANDIADGGVRAISQLDTCAQLKRSDPPSPPFLQQQQQVPVLHQNE<br>AVSALSFSRYAPFDDQSLILPQRTQRVSVLRQAVHRPAADTTSGISDF<br>ELPYTLPSPKKTPLCERKGPLVVKEPYAKAHGPVYVVKREDSKMYEMN<br>ASVVSRGAMASLSRSFRCGWCTGGRLVAPMFAWVRDGTESRHSVDEVL<br>GSRVVMSTVYFAHATSKHYLQSCAISVLRALCRHAHRVDASSDKEGFF<br>PLLELGLCRDTGSTSLSTEKLREVIAAVDAVRFDGASAVGESTARQAK<br>TILSLLDALYGLPDADEADKNAITERRYLTQLRRRNLNSWLKTELEYM<br>DLWTDLDVDANPSQKLLRKLLCGKLREASSVAKALGSTELSRVVGICG<br>EGNHFGSYVQTSDNSCIDEALGIRERVVSLLSGIVEPFVSQPQYAWAE<br>DEKGGRTVAKVPLAATWKQLLGVFAFYGCTPDSSAEETIDCFLKRLRA<br>PTSRRENSFPPYAERISADKLGTGRGRSFVSLGEWFPDAALSLLEGFA<br>MGVAPAATALHPHSSSYCATDYLTPFIIIVSVRALKLQRTDTYDDAET<br>KALLGFAAALECLSDAWFWALLPLHMIVDDKCRAVAVEQCLRRNAHRF<br>QGGACRTNTDYVHLAELLKINARLLDVEMLLEKVPADAPVNAPSIRTH<br>SSLQEALHRFSRGFTKR |
| Q64K55 | Q64K55_ARGTR | SSALFNAGVLNASNIDTLGSRVLSALLNGVSSAAQGLGINVDSGSVQS<br>DISSSSSFLSTSSSSASYSQASASSTSGAGYTGPSGPSTGPSGYPGPL<br>GGGAPFGQSGFGGSDGPQGGFGATGGASAGLISRVANALANTSTLRTV<br>LRTGVSQQIASSVVQRAAQSLASTLGVDGNNLARFAVQAVSRLPAGSD<br>TSAYAQAFSSALFNAGVLNASNIDTLGSRVLSALLNGVSSAAQGLGIN<br>VDSGSVQSDISSSSSFLSTSSSSASYSQASASSTSGAGYTGPSGPSTG<br>PSGYPGLLGGGAPFGQSGFGGSDGPQGGFGATGGASAGLISRVANALA<br>NTSTLRTVLRTGVSQQIASSVVQRAAQSLASTLGVDGNNLARFAVQAV<br>SRLPAGSDTSAYAQAFSSALFNAGVLNASNIDTLGSRVLSALLNGVSS<br>AAQGLGINVDSGSVQSDISSSSSFLSTSSSSASYSQASASSTSGAGYT<br>GPSGPSTGPSGYPGPLGGGAPFGQSGFGGSDGPQGGFGATGGASAGLI<br>SRVANALANTSTLRTVLRTGVSQQIASSVVQRAAQSLASTLGVDGNNL<br>ARFAVQAVSRLPAGSDTSAYAQAFSSALFNAGVLNASNIDTLGSRVLS<br>ALLNGVSSAAQGLGINVDSGSVQSDISSSSSFLSTSSSSASYSQASAS<br>STSGAGYTGPSGPSTGPSGYPGPLGGGAPFGQSGFGGSAGPQGGFGAT<br>GGASAGLISRVANALANTSTLRTVLRTGVSQQIASSVVQRAAQSLAST<br>LGVDGNNLARFAVQAVSRLPAGSDTSAYAQAFSSALFNAGVLNASNID<br>TLGSRVLSALLNGVSSAAQGLGINVDSGSVQSDISSSSSFLSTSSSSA<br>SYSQASASSTSGAGYTGPSGPSTGPSGYPGPLGGGAPFGQSGFGGSAG<br>PQGGFGATGGASAGLISRVANALANTSTLRTVLRTGVSQQIASSVVQR<br>AAQSLASTLGVDGNNLARFAVQAVSRLPAGSDTSAYAQAFSSALFNAG<br>VLNASNIDTLGSRVLSALLNGVSSAAQGLGINVDSGSVQSDISSSSSF<br>LSTSSSSASYSQASASSTSGTGYTGPSGPSTGPSGYPGPLGGGAPFGQ<br>SGFGGSAGPQGGFGATGGASAGLISRVANALANTSTLRTVLRTGVSQQ<br>IASSVVQRAAQSLASTLGVDGNNLARFAVQAVSRLPAGSDTSAYAQAF<br>SSALFNAGVLNASNIDTLGSRVLSALLNGVSSAAQGLGINVDSGSVQS<br>DISSSSSFLSTSSSSASYSQASASSTSGAGYTGPSGPSTGPSGYPGPL<br>GGGAPFGQSGFGGSAGPQGGFGATGGASAGLISRVANALANTSTLRTV<br>LRTGVSQQIASSVVQRAAQSLASTLGVDGNNLARFAVQAVSRLPAGSD |

| | | |
|---|---|---|
| | | TSAYAQAFSSALFNAGVLNASNIDTLGSRVLSALLNGVSSAAQGLGIN<br>VDSGSVQSDISSSSSFLSTSSSSASYSQASASSTSGAGYTGPSGPSTG<br>PSGYPGPLGGGAPFGQSGFGGSDGPQGGFGATGGASAGLISRVANALA<br>NTSTLRTVLRTGVSQQIASSVVQRAAQSLASTLGVDGNNLARFAVQAV<br>SRLPAGSDTSAYAQAFSSALFNAGVLNASNIDTLGSRVLSALLNGVSS<br>AAQGLGINVDSGSVQSDISSSSSFLSTSSSSASYSQASASSTSGAGYT<br>GPSGPSTGPSGYPGPLGGGAPFGQSGFGGSAGPQGGFGATGGASAGLI<br>SRVANALANTSTLRTVLRTGVSQQIASSVVQRAAQSLASTLGVDGNNL<br>ARFAVQAVSRLPAGSDTSAYAQAFSSALFNAGVLNASNIDTLGSRVLS<br>ALLNGVSSAAQGLGINVDSGSVQSDISSSSSFLSTSSSSASYSQASAS<br>STSGAGYTGPSGPSTGPSGYPGPLGGGAPFGQSGFGGSAGPQGGFGAT<br>GGASAGLISRVANALANTSTLRTVLRTGVSQQIASSVVQRAAQSLAST<br>LGVDGNNLARFAVQAVSRLPAGSDTSAYAQAFSSALFNAGVLNASNID<br>TLGSRVLSALLNGVSSAAQGLGINVDSGSVQSDISSSSSFLSTSSSSA<br>SYSQASASSTSGAGYTGPSGPSTGPSGYPGPLGGGAPFGQSGFGGSAG<br>PQGGFGATGGASAGLISRVANALANTSTLRTVLRTGVSQQIASSVVQR<br>AAQSLASTLGVDGNNLARFAVQAVSRLPAGSDTSAYAQAFSSALFNAG<br>VLNASNIDTLGSRVLSALLNGVSSAAQGLGINVDSGSVQSDISSSSSF<br>LSTSSSSASYSQASASSTSGAGYTGPSGPSTGPSGYPGPLGGGAPFGQ<br>SGFGGSAGPQGGFGATGGASAGLISRVANALANTSTLRTVLRTGVSQQ<br>IASSVVQRAAQSLASTLGVDGNNLARFAVQAVSRLPAGSDTSAYAQAF<br>SSALFNAGVLNASNIDTLGSRVLSALLNGVSSAAQGLGINVDSGSVQS<br>DISSSSSFLSTSSSSASYSQALASSTSGAGYTGPSGPSTGPSGYPGPL<br>GGGAPFGQSGFGGSAGPQGGFGATGGASAGLISRVANALANTSTLRTV<br>LRTGVSQQIASSVVQRAAQSLASTLGVDGNNLARFAVQAVSRLPAGSD<br>TSAYAQAFSSALFNAGVLNASNIDTLGSRVLSALLNGVSSAAQGLGIN<br>VDSGSVQSDISSSSSFLSTSSSSASYSQASASSTSGAGYTGPSGPSTG<br>PSGYPGPLSGGASFGSGQSSFGQTSAFSASGAGQSAGVSVISSLNSPV<br>GLRSASAASRLSQLTSSITNAVGANGVDANSLARSLQSSFSALRSSGM<br>SSSDAKIEVLLETIVGLLQLLSNTQVRGVNPATASSVANSAARSFELV<br>LA |
| Q02629 | NU100_YEAST | MFGNNRPMFGGSNLSFGSNTSSFGGQQSQQPNSLFGNSNNNNNSTSNN<br>AQSGFGGFTSAAGSNSNSLFGNNNTQNNGAFGQSMGATQNSPFGSLNS<br>SNASNGNTFGGSSSMGSFGGNTNNAFNNNSNSTNSPFGFNKPNTGGTL<br>FGSQNNNSAGTSSLFGGQSTSTTGTFGNTGSSFGTGLNGNGSNIFGAG<br>NNSQSNTTGSLFGNQQSSAFGTNNQQGSLFGQQSQNTNNAFGNQNQLG<br>GSSFGSKPVGSGSLFGQSNNTLGNTTNNRNGLFGQMNSSNQGSSNSGL<br>FGQNSMNSSTQGVFGQNNNQMQINGNNNNSLFGKANTFSNSASGGLFG<br>QNNQQQGSGLFGQNSQTSGSSGLFGQNNQKQPNTFTQSNTGIGLFGQN<br>NNQQQQSTGLFGAKPAGTTGSLFGGNSSTQPNSLFGTTNVPTSNTQSQ<br>QGNSLFGATKLTNMPFGGNPTANQSGSGNSLFGTKPASTTGSLFGNNT<br>ASTTVPSTNGLFGNNANNSTSTTNTGLFGAKPDSQSKPALGGGLFGNS<br>NSNSSTIGQNKPVFGGTTQNTGLFGATGTNSSAVGSTGKLFGQNNNTL<br>NVGTQNVPPVNNTTQNALLGTTAVPSLQQAPVTNEQLFSKISIPNSIT<br>NPVKATTSKVNADMKRNSSLTSAYRLAPKPLFAPSSNGDAKFQKWGKT<br>LERSDRGSSTSNSITDPESSYLNSNDLLFDPDRRYLKHLVIKNNKNLN<br>VINHNDDEASKVKLVTFTTESASKDDQASSSIAASKLTEKAHSPQTDL<br>KDDHDESTPDPQSKSPNGSTSIPMIENEKISSKVPGLLSNDVTFFKNN<br>YYISPSIETLGNKSLIELRKINNLVIGHRNYGKVEFLEPVDLLNTPLD<br>TLCGDLVTFGPKSCSIYENCSIKPEKGEGINVRCRVTLYSCFPIDKET<br>RKPIKNITHPLLKRSIAKLKENPVYKFESYDPVTGTYSYTIDHPVLT |
| Q02630 | NU116_YEAST | MFGVSRGAFPSATTQPFGSTGSTFGGQQQQQQPVANTSAFGLSQQTNT<br>TQAPAFGNFGNQTSNSPFGMSGSTTANGTPFGQSQLTNNNASGSIFGG<br>MGNNTALSAGSASVVPNSTAGTSIKPFTTFEEKDPTTGVINVFQSITC<br>MPEYRNFSFEELRFQDYQAGRKFGTSQNGTGTTFNNPQGTTNTGFGIM<br>GNNNSTTSATTGGLFGQKPATGMFGTGTGSGGGFGSGATNSTGLFGSS<br>TNLSGNSAFGANKPATSGGLFGNTTNNPTNGTNNTGLFGQQNSNTNGG<br>LFGQQQNSFGANNVSNGGAFGQVNRGAFPQQQTQQGSGGIFGQSNANA<br>NGGAFGQQQGTGALFGAKPASGGLFGQSAGSKAFGMNTNPTGTTGGLF<br>GQTNQQQSGGGLFGQQQNSNAGGLFGQNNQSQNQSGLFGQQNSSNAFG<br>QPQQQGGLFGSKPAGGLFGQQQGASTFASGNAQNNSIFGQNNQQQQST<br>GGLFGQQNNQSQSQPGGLFGQTNQNNNQPFGQNGLQQPQQNNSLFGAK<br>PTGFGNTSLFSNSTTNQSNGISGNNLQQQSGGLFQNKQQPASGGLFGS<br>KPSNTVGGGLFGNNQVANQNNPASTSGGLFGSKPATGSLFGGTNSTAP<br>NASSGGIFGSNNASNTAATTNSTGLFGNKPVGAGASTSAGGLFGNNNN |

| | | |
|---|---|---|
| | | SSLNNSNGSTGLFGSNNTSQSTNAGGLFQNNTSTNTSGGGLFSQPSQS<br>MAQSQNALQQQQQQQRLQIQNNNPYGTNELFSKATVTNTVSYPIQPSA<br>TKIKADERKKASLTNAYKMIPKTLFTAKLKTNNSVMDKAQIKVDPKLS<br>ISIDKKNNQIAISNQQEENLDESILKASELLFNPDKRSFKNLINNRKM<br>LIASEEKNNGSQNNDMNFKSKSEEQETILGKPKMDEKETANGGERMVL<br>SSKNDGEDSATKHHSRNMDEENKENVADLQKQEYSEDDKKAVFADVAE<br>KDASFINENYYISPSLDTLSSYSLLQLRKVPHLVVGHKSYGKIEFLEP<br>VDLAGIPLTSLGGVIITFEPKTCIIYANLPNRPKRGEGINVRARITCF<br>NCYPVDKSTRKPIKDPNHQLVKRHIERLKKNPNSKFESYDADSGTYVF<br>IVNHAAEQT |
| F4ID16 | NU98B_ARATH | MFGSSNNNPFGQSSISSPFGTQTHSLFGQTNNNASNNPFATKPFGTST<br>PFGAQTGSSMFGGTSTGVFGAPQTSSPFGASPQAFGSSTQAFGASSTP<br>SFGSSNSPFGGTSTFGQKSFGLSTPQSSPFGSTTQQSQPAFGNSTFGS<br>STPFGASTTPAFGASSTPAFGVSNTSGFGATNTPGFGATNTTGFGGSS<br>TPGFGASSTPAFGSTNTPAFGASSTPLFGSSSSPAFGASPAPAFGSSG<br>NAFGNNTFSSGGAFGSSSTPTFGASNTSAFGASSSPSFNFGSSPAFGQ<br>STSAFGSSSFGSTQSSLGSTPSPFGAQGAQASTSTFGGQSTIGGQQGG<br>SRVIPYAPTTDTASGTESKSERLQSISAMPAHKGKNMEELRWEDYQRG<br>DKGGQRSTGQSPEGAGFGVTNSQPSIFSTSPAFSQTPVNPTNPFSQTT<br>PTSNTNFSPSFSQPTTPSFGQPTTPSFRSTVSNTTSVFGSSSSLTTNT<br>SQPLGSSIFGSTPAHGSTPGFSIGGFNNSQSSPLFGSNPSFAQNTTPA<br>FSQTSPLFGQNTTPALGQSSSVFGQNTNPALVQSNTFSTPSTGFGNTF<br>SSSSSLTTSISPFGQITPAVTPFQSAQPTQPLGAFGFNNFGQTQIANT<br>TDIAGAMGTFSQGNFKQQPALGNSAVMQPTPVTNPFGTLPALPQISIA<br>QGGNSPSIQYGISSMPVVDKPAPVRVSPLLTSRHLLQRRVRLPTRKYR<br>PSDDGPKVPFFSDEEENSSTPKADAFFIPRENPRALFIRPVERVKSEH<br>PKDSPTPLQENGKRSNGVTNGANHETKDNGAIREAPPVKVNQKQNGTH<br>ENHGGDKNGSHSSPSGADIESLMPKLHHSEYFTEPRIQELAAKERVEQ<br>GYCKRVKDFVVGRHGYGSIKFLGETDVCRLDLEMVVQFKNREVNVYMD<br>ESKKPPVGQGLNKPAVVTLLNIKCMDKKTGTQVMEGERLDKYKEMLKR<br>KAGEQGAQFVSYDPVNGEWTFKVEHFSSYKLGDEYDV |
| Q54EQ8 | NUP98_DICDI | MFGGQFGSFGAKPAATASPFGAPSAAPTTSLFGSTAPSSGFGGFGSTA<br>QTTQPTTGGFGGFGGFGGATTTQQPAASPFGGGGTGGSGLFGSSAQTT<br>TQQPGASPFGGGFGTTTTTTTQQPGASPFGGTGGGLFGSSAQTTTQQQ<br>GASPFGGFGGATTTQPSLLSGATGGFGGFGGSTTSGTQLGGGGGATSG<br>AFGGSSSPFGGSGGATTSSPFGGGGGSFGATTQKQYGTPIPYQQTTIE<br>GNTFVSISAMPQYNDRSFEELRFEDITHRKDIVYKTGGGSGGGNSLFG<br>STPTTQPSSPFGAQTTTQTTGGLFGGQTTTSPFGGQTSATPGSSLFGS<br>TQPTQQQTSGGLFGSVQPTQQQAGGGLFGSMPSTGGSSLFGSTQPTQQ<br>QTGGAQPTQSLFGGQTQTTTSPFGSQTSTPFGQPQQTNTGSGLFGAQQ<br>TQQTNTGGGLFGAQPTQQTSGGGLFGTQPTSGTGLFGTSPTAGGTGLF<br>GTTQPTSQGTGLFGTTQPTTQGTGLFGTSPTSGTGLFGSTPTSGTGLF<br>GSTPTSGTGLFGSAQPPQNQQSQTSLFGNTGTGATNTGTGLFGSAQPS<br>SNPGGGLFGSAQPSTTTGGLFGSNQPTAQPTTSLFGNTTGSVGGLGAT<br>PNITSGLFGSNPAQTGGLFGSTQPTTQTSLFGNTGSTGGLGAQNGGGL<br>FGNLSQPTATAGQGLSGGLFGNLSQPTATAGQGLSSGGLFGNTLLGQP<br>STQGLSSALPTLGLGLMGGQPQQTQQLPQGSLMLQQTQQPLQQQPLQQ<br>QQQPLQQSTIQLNNQINSASPYFPISSPAPFATFVKDLTSTSKVVSPP<br>SYTQRSLSHHGYIPKSTTKLVPRRGPNNVDLGFSVIQNQNGLFPIDKF<br>ITKHSKSLNINTTNETEDTLRSLNTKSSSLFNNNNNNNNNNNNNRNVNT<br>NVNDYQNNGLPSSSLYNSNINQLSNNNNNNNNIYNNNNNNNINNNNNN<br>NNISTQFNLRNNQSSSDNLNNDKSLSSSSSNKSQQQQQKEQKEEQPPK<br>PIKEKEFINPNAPKLTRDGYQCVPSIKELSKKTDKELSSVQGFTISRD<br>GCGSIYFPGSTNLVALDLDDIVDIEPREVSVYKDEETKPEIGYGLNRD<br>AVVTLENCWPKNKNGEVVKEDGTILDKYENALKKVSAKSDCGFVSYSR<br>SNGTWVFTVKHFSKYSAPDFDEDDQQMQQQTQQKQQQTQPSKVTFQQP<br>STKLTKPKFTANLDNFDSDSETSSGDENQDEMVPQKKTPFIKRVSNRE<br>SGLFDTPSVVPMSEKIETTPSKIARVSEPTSQSSRMSNNALKFSTFNP<br>QQQQLQSSRFKSTGLSILSNPVKNLIQSDVNNEQSMFSNTTTTSTTRI<br>QPLPSQQHLVQPIPTTISLNKNYFSKVRIDPQVYDRIVPKEESITNQY<br>RMKNERLNHSTQDVSLFMRRSFRVGWAPGGKLISITKSSFKNLLIKKL<br>PTDTKEDKKESIIKFLKNHHSHSSLVPENLKSIGWFSISNVQEQIESQ<br>LTLNVPSSQSVYYNRIWSLISNLWGNVLKGNGSKYINTNYSEDTIRKL<br>NLNQWLKDVIAPLLRDEMDSLRKKTNSNYLEQIFSYLSAKQIKEASDL<br>ANENKDFRLATMMSQIWSSSESGKELILKQLTTYHSNGSDEFINEKRL |

| | | |
|---|---|---|
| | | EILHLIAGSVNKIYKNLNDWIRCFAVSFWFKYSLEYSIEDSVENFERS<br>FNAHRSVYPLPPYLIKSTSTNSKQIEEQQHYYDICFLLLKLFAVNRGS<br>SHFDKFKNIFYPENIGQDLLDYHLSWNLYTVLKSIPSLNKQPDLVNAS<br>NLHSSFALQLERLGLWQWSIYVLLHTPDQSNHVREEAVKSLIARAAPV<br>ITSEDRVFLTTKLHIPEIWIDEAKAWYSGYDCNNDIYDQIDALFKSYQ<br>YTKIHDIIFSNIGPNYIIQKRYHSLKDLLIRLEPHSSFISTWRYGGSI<br>FLEFADICIQYKEILSQLSNTAEEIQRTKYYVNLKDITTRIVNILSDI<br>SKITQSSEIKNTSASYKQSLSFMSEALITKASLLRDLPESIVKLVSTN<br>NLVSTLNSLPLTQDYRSKNLESLTDQIQDTLLNSIYQ |
| Q13148 | TADBP_HUMAN | MSEYIRVTEDENDEPIEIPSEDDGTVLLSTVTAQFPGACGLRYRNPVS<br>QCMRGVRLVEGILHAPDAGWGNLVYVVNYPKDNKRKMDETDASSAVKV<br>KRAVQKTSDLIVLGLPWKTTEQDLKEYFSTFGEVLMVQVKKDLKTGHS<br>KGFGFVRFTEYETQVKVMSQRHMIDGRWCDCKLPNSKQSQDEPLRSRK<br>VFVGRCTEDMTEDELREFFSQYGDVMDVFIPKPFRAFAFVTFADDQIA<br>QSLCGEDLIIKGISVHISNAEPKHNSNRQLERSGRFGGNPGGFGNQGG<br>FGNSRGGGAGLGNNQGSNMGGGMNFGAFSINPAMMAAAQAALQSSWGM<br>MGMLASQQNQSGPSGNNQNQGNMQREPNQAFGSGNNSYSGSNSGAAIG<br>WGSASNAGSGSGFNGGFGSSMDSKSSGWGM |
| Q9PVZ2 | Q9PVZ2_XENLA | MEDDTDLPPERETKDFQFRQLKKVRLFDYPADLPKQRSNLLVISNKYG<br>LLFVGGFMGLKVFHTKDILVTVKPKENANKTVVGPQGIHVPMNSPIHH<br>LALSSDNLTLSVCMTSAEQGSSVSFYDVRTLLNESKQNKMPFASCKLL<br>RDPSSSVTDLQWNPTLPSMVAVCLSDGSISVLQVTDTVSVFANLPATL<br>GVTSVCWSPKGKQLAVGKQNGTVVQYLPSLQEKKVIPCPSFYDSDNPV<br>KVLDVLWLSTYVFTVVYAAADGSLEASPQLVIVTLPKKEDKRAERFLN<br>FTETCYSICSERQHHFFLNYIEDWEILLAASAASVDVGVIARPPDQVG<br>WEQWLLEDSSRAEMPMTENNDDTLPMGVALDYTCQLEVFISESQILPP<br>VPVLLLLSTDGVLCPFHVVNLNQGVKPLTTSPEQLSLDGEREMKVVGG<br>TAVSTPPAPLTSVSAPAPPASAAPRSAAPPPYPFGLSTASSGAPTPVL<br>NPPASLAPAATPTKTTSQPAAAATSIFQPAGPAAGSLQPPSLPAFSFS<br>SANNAANASAPSSFPFGAAMVSSNTAKVSAPPAMSFQPAMGTRPFSLA<br>TPVTVQAATAPGFTPTPSTVKVNLKDKFNASDTPPPATISSAAALSFT<br>PTSKPNATVPVKSQPTVIPSQASVQPNRPFAVEAPQAPSSVSIASVQK<br>TVRVNPPATKITPQPQRSVALENQAKVTKESDSILNGIREEIAHFQKE<br>LDDLKARTSRACFQVGSEEEKRQLRTESDGLHSFFLEIKETTESLRGE<br>FSAMKIKNLEGFASIEDVQQRNKLKQDPKYLQLLYKKPLDPKSETQMQ<br>EIRRLNQYVKNAVQDVNDVLDLEWDQYLEEKQKKKGIIIPERETLFNS<br>LANHQEIINQQRPKLEQLVENLQKLRLYNQISQWNVPDSSTKSFDVEL<br>ENMQKTLSQTAIDTQTKPQAKLPAKISPVKQSQLRNFLSKRKTPPVRS<br>LAPANLSRSAFLAPSFFEDLDDVSSTSSLSDMADNDNRNPPPKEIERQ<br>ETPPPESTPVRVPKHAPVARTTSVQPGLGTASLPFQSGLHPATSTPVA<br>PSQSIRVIPQGADSTMLATKTVKHGAPNITAAQKAAVAAMRRQTASQI<br>PAASLTESTLQTVPQVVNVKELKNNGPGPTIPTVIGPTVPQSAAQVIH<br>QVLATVGSVSARQAAPAAPLKNPPASASSIAPQTWQGSAPNKPAAQAI<br>PKSDPSASQAPAPSVSQVNKPVSFSPAAGGFSFSNVTSAPVTSALGSS<br>SAGCAATARDSNQASSYMFGGTGKSLGSEGSFSFASLKPASSSSSSSV<br>VEPTMSKPSVVTAASTTATVTSTTAASSKPGEGLFQGFSGGETLGSFS<br>GLRVGQADEASKVEVAKTPTAAQPVKLPSNPVLFSFAGAPQPAKVGEA<br>PSTTSSTSASLFGNVQLASAGSTASAFTQSGSKPAFTFGIPQSTSTTA<br>GASSAIPASFQSLLVSAAPATTTPSAPINSGLDVKQPIKPLSEPADSS<br>SSQQQTLTTQSAAEQVPTVTPAATTATALPPPVPTIPSTAEAKIEGAA<br>APAIPASVISSQTVPFTSTVLASQTPLASTPAGGPTSQVPVLVTTAPP<br>VTTESAQTVSLTGQPVAGSSAFAQSTVTAASTPVFGQALASGAAPSPF<br>AQPTSSSVSTSANSSTGFGTSAFGATGGNGGFGQPSFGQAPLWKGPAT<br>SQSTLPFSQPTFGTQPAFGQPAASTATSSAGSLFGCTSSASSFSFGQA<br>SNTSGTSTSGVLFGQSSAPVFGQSAAFPQAAPAFGSASVSTTTTASFG<br>FGQPAGFASGTSGSLFNPSQSGSTSVFGQQPASSSGGLFGAGSGGAST<br>VGLFSGLGAKPSQEAANKNPFGSPGSSGFGSAGASNSSNLFGNSGAKA<br>FGFGGTSFGDKPSATFSAGGSVASQGFSFNSPTKTGGFGAAPVFGSPP<br>TFGGSPGFGGSPAFGTAAAFSNTLGSTGGKVFGEGTSAATTGGFGFGS<br>NSSTAAFGSLATQNTPTFGSISQQSPGFGGQSSGFSGFGAGPGAAAGN<br>TGGFGFGVSNPTSPGFGCWRS |
| G5EEH9 | NUP98_CAEEL | MFGQNKSFGSSSFGGGSSGSGLFGQNNQNNQNKGLFGQPANNSGTTGL<br>FGAAQNKPAGSIFGAASNTSSIFGSPQQPQNNQSSLFGGGQNNANRSI<br>FGSTSSAAPASSSLFGNNANNTGTSSIFGSNNNAPSGGGLFGASTVSG |

| | | |
|---|---|---|
| | | TTVKFEPPISSDTMMRNGTTQTISTKHMCISAMSKYDGKSIEELRVED<br>YIANRKAPGTGTTSTGGGLFGASNTTNQAGSSGLFGSSNAQQKTSLFG<br>GASTSSPFGGNTSTANTGSSLFGNNNANTSAASGSLFGAKPAGSSLFG<br>STATTGASTFGQTTGSSLFGNQQPQTNTGGSLFGNTQNQNQSGSLFGN<br>TGTTGTGLFGQAQQQPQQQSSGFSFGGAPAATNAFGQPAAANTGGSLF<br>GNTSTANTGSSLFGAKPATSTGFTFGATQPTTTNAFGSTNTGGGLFGN<br>NAAKPGGLFGNTTNTGTGGGLFGSQPQASSGGLFGSNTQATQPLNTGF<br>GNLAQPQIVMQQQVAPVPVIGVTADVLQMQANMKSLKSQLTNAPYGDS<br>PLLKYNANPEIDGKSSPASTQRQLRFLAAKKGALSSSSDAQDSSFIIP<br>PISKVMSDLSPAVTRSADVTKDLNYTSKEAPPSLARGLRNSTFNPNMS<br>LTNRSVHESSALDKTIDSALDASMNGTSNRLGVRGSVRRSNLKQLDMS<br>LLADSSRVGRESRVADPDALPRISESERRQDVVTSTPAVDPVQAVIQR<br>HNDRNRDPPSLNLDTTCDEHTGLEPVSAATSSAASVVSTPSEETVNVN<br>SAAGVKLTKPDYFSLPTINEMKNMIKNGRVVLEDGLTVGRSSYGSVYW<br>PGRVELKDVALDEIVVFRHREVTVYPNEEEKAPEGQELNRPAEVTLER<br>VWYTDKKTKKEVRDVVKLSEIGWREHLERQTIRMGAAFKDFRAETGSW<br>VFRVDHFSKYGLADDDEPMDGSPPQQALQASSPLQVIDMNTSARDVNN<br>QVQRKKVHKATDAHHQEIILERVPAPAALGDVVPIIRRVNRKGLGGGT<br>LDDSREESCIGNMTTEFNESGHDSIIEEGQQPEKKPKLELLADLEYES<br>SRFIRNLQELKVMPKANDPAHRFHGGGHSAKMIGYGKSKLIDIGIVKG<br>RSSHVGWSETGCLVWSAQPRHNQVLFGTIDRTSDVNENTLISMLDVNV<br>HVSETSRKGPSSQSNSVKSSLTSNFVTYSDSYSSMFAKYIDVAQAGGY<br>DGHVSVWKLISALFPYERREGWSFERGEEIGEWLRTEAVKSVPDDRSA<br>DTSSNGVWNQLCLGDIDKAFQIAIDNNQPQLATMLQTSAVCPEATVHC<br>FKAQLDNWKKCETLHLIPKETLKCYVLMSGLSHYEWDQDGKNHSINCL<br>DGLNWIQALGLHVWYLRAWTGLEESYDAYQKDVNAGRAASNRGDLPGE<br>LIKLACESQHSVEVVLDCAAGENPNDYFLQWHVWSLLYSVGYRTMSKT<br>SETRLHRNYSSQLEASSLSKYALFVLQHIDDDEERSTAVRSLLDRIAR<br>FTDNDMFDSISEQFDIPSEWIADAQFSIAKSVDDSTQLFELAVAAKNY<br>LEICRLFVDDIAPTAVVAGDHDALKAACAMVRPFENQIPEWGATGMVY<br>TDYCRLINLIENDAEEELLQDVLESLETRLHAPTISKNSLQKLSLQTI<br>GRVLFEYRADKNTLPEWTKLLGHRQMFKIFRDRSSWGIERFTIEFD |
| P52948 | NUP98_HUMAN | MFNKSFGTPFGGGTGGFGTTSTFGQNTGFGTTSGGAFGTSAFGSSNNT<br>GGLFGNSQTKPGGLFGTSSFSQPATSTSTGFGFGTSTGTANTLFGTAS<br>TGTSLFSSQNNAFAQNKPTGFGNFGTSTSSGGLFGTTNTTSNPFGSTS<br>GSLFGPSSFTAAPTGTTIKFNPPTGTDTMVKAGVSTNISTKHQCITAM<br>KEYESKSLEELRLEDYQANRKGPQNQVGAGTTTGLFGSSPATSSATGL<br>FSSSTTNSGFAYGQNKTAFGTSTTGFGTNPGGLFGQQNQQTTSLFSKP<br>FGQATTTQNTGFSFGNTSTIGQPSTNTMGLFGVTQASQPGGLFGTATN<br>TSTGTAFGTGTGLFGQTNTGFGAVGSTLFGNNKLTTFGSSTTSAPSFG<br>TTSGGLFGNKPTLTLGTNTNTSNFGFGTNTSGNSIFGSKPAPGTLGTG<br>LGAGFGTALGAGQASLFGNNQPKIGGPLGTGAFGAPGFNTTTATLGFG<br>APQAPVALTDPNASAAQQAVLQQHINSLTYSPFGDSPLFRNPMSDPKK<br>KEERLKPTNPAAQKALTTPTHYKLTPRPATRVRPKALQTTGTAKSHLF<br>DGLDDDEPSLANGAFMPKKSIKKLVLKNLNNSNLFSPVNRDSENLASP<br>SEYPENGERFSFLSKPVDENHQQDGDEDSLVSHFYTNPIAKPIPQTPE<br>SAGNKHSNSNSVDDTIVALNMRAALRNGLEGSSEETSFHDESLQDDRE<br>EIENNSYHMHPAGIILTKVGYYTIPSMDDLAKITNEKGECIVSDFTIG<br>RKGYGSIYFEGDVNLTNLNLDDIVHIRRKEVVVYLDDNQKPPVGEGLN<br>RKAEVTLDGVWPTDKTSRCLIKSPDRLADINYEGRLEAVSRKQGAQFK<br>EYRPETGSWVFKVSHFSKYGLQDSDEEEEEHPSKTSTKKLKTAPLPPA<br>SQTTPLQMALNGKPAPPPQSQSPEVEQLGRVVELDSDMVDITQEPVLD<br>TMLEESMPEDQEPVSASTHIASSLGINPHVLQIMKASLLTDEEDVDMA<br>LDQRFSRLPSKADTSQEICSPRLPISASHSSKTRSLVGGLLQSKFTSG<br>AFLSPSVSVQECRTPRAASLMNIPSTSSWSVPPPLTSVFTMPSPAPEV<br>PLKTVGTRRQLGLVPREKSVTYGKGKLLMDMALFMGRSFRVGWGPNWT<br>LANSGEQLNGSHELENHQIADSMEFGFLPNPVAVKPLTESPFKVHLEK<br>LSLRQRKPDEDMKLYQTPLELKLKHSTVHVDELCPLIVPNLGVAVIHD<br>YADWVKEASGDLPEAQIVKHWSLTWTLCEALWGHLKELDSQLNEPREY<br>IQILERRRAFSRWLSCTATPQIEEEVSLTQKNSPVEAVFSYLTGKRIS<br>EACSLAQQSGDHRLALLLSQFVGSQSVRELLTMQLVDWHQLQADSFIQ<br>DERLRIFALLAGKPVWQLSEKKQINVCSQLDWKRSLAIHLWYLLPPTA<br>SISRALSMYEEAFQNTSDSDRYACSPLPSYLEGSGCVIAEEQNSQTPL<br>RDVCFHLLKLYSDRHYDLNQLLEPRSITADPLDYRLSWHLWEVLRALN<br>YTHLSAQCEGVLQASYAGQLESEGLWEWAIFVLLHIDNSGIREKAVRE |

| | | |
|---|---|---|
| | | LLTRHCQLLETPESWAKETFLTQKLRVPAKWIHEAKAVRAHMESDKHL<br>EALCLFKAEHWNRCHKLIIRHLASDAIINENYDYLKGFLEDLAPPERS<br>SLIQDWETSGLVYLDYIRVIEMLRHIQQVDCSGNDLEQLHIKVTSLCS<br>RIEQIQCYSAKDRLAQSDMAKRVANLLRVVLSLHHPPDRTSDSTPDPQ<br>RVPLRLLAPHIGRLPMPEDYAMDELRSLTQSYLRELAVGSL |
| P49790 | NU153_HUMAN | MASGAGGVGGGGGGKIRTRRCHQGPIKPYQQGRQQHQGILSRVTESVK<br>NIVPGWLQRYFNKNEDVCSCSTDTSEVPRWPENKEDHLVYADEESSNI<br>TDGRITPEPAVSNTEEPSTTSTASNYPDVLTRPSLHRSHLNFSMLESP<br>ALHCQPSTSSAFPIGSSGFSLVKEIKDSTSQHDDDNISTTSGFSSRAS<br>DKDITVSKNTSLPPLWSPEAERSHSLSQHTATSSKKPAFNLSAFGTLS<br>PSLGNSSILKTSQLGDSPFYPGKTTYGGAAAAVRQSKLRNTPYQAPVR<br>RQMKAKQLSAQSYGVTSSTARRILQSLEKMSSPLADAKRIPSIVSSPL<br>NSPLDRSGIDITDFQAKREKVDSQYPPVQRLMTPKPVSIATNRSVYFK<br>PSLTPSGEFRKTNQRIDNKCSTGYEKNMTPGQNREQRESGFSYPNFSL<br>PAANGLSSGVGGGGGKMRRERTRFVASKPLEEEEMEVPVLPKISLPIT<br>SSSLPTFNFSSPEITTSSPSPINSSQALTNKVQMTSPSSTGSPMFKFS<br>SPIVKSTEANVLPPSSIGFTFSVPVAKTAELSGSSSTLEPIISSSAHH<br>VTTVNSTNCKKTPPEDCEGPFRPAEILKEGSVLDILKSPGFASPKIDS<br>VAAQPTATSPVVYTRPAISSFSSSGIGFGESLKAGSSWQCDTCLLQNK<br>VTDNKCIACQAAKLSPRDTAKQTGIETPNKSGKTTLSASGTGFGDKFK<br>PVIGTWDCDTCLVQNKPEAIKCVACETPKPGTCVKRALTLTVVSESAE<br>TMTASSSSCTVTTGTLGFGDKFKRPIGSWECSVCCVSNNAEDNKCVSC<br>MSEKPGSSVPASSSSTVPVSLPSGGSLGLEKFKKPEGSWDCELCLVQN<br>KADSTKCLACESAKPGTKSGFKGFDTSSSSSNSAASSSFKFGVSSSSS<br>GPSQTLTSTGNFKFGDQGGFKIGVSSDSGSINPMSEGFKFSKPIGDFK<br>FGVSSESKPEEVKKDSKNDNFKFGLSSGLSNPVSLTPFQFGVSNLGQE<br>EKKEELPKSSSAGFSFGTGVINSTPAPANTIVTSENKSSFNLGTIETK<br>SASVAPFTCKTSEAKKEEMPATKGGFSFGNVEPASLPSASVFVLGRTE<br>EKQQEPVTSTSLVFGKKADNEEPKCQPVFSFGNSEQTKDENSSKSTFS<br>FSMTKPSEKESEQPAKATFAFGAQTSTTADQGAAKPVFSFLNNSSSSS<br>STPATSAGGGIFGSSTSSSNPPVATFVFGQSSNPVSSSAFGNTAESST<br>SQSLLFSQDSKLATTSSTGTAVTPFVFGPGASSNNTTTSGFGFGATTT<br>SSSAGSSFVFGTGPSAPSASPAFGANQTPTFGQSQGASQPNPPGFGSI<br>SSSTALFPTGSQPAPPTFGTVSSSSQPPVFGQQPSQSAFGSGTTPNSS<br>SAFQFGSSTTNFNFTNNSPSGVFTFGANSSTPAASAQPSGSGGFPFNQ<br>SPAAFTVGSNGKNVFSSSGTSFSGRKIKTAVRRRK |
| Q5EAX5 | Q5EAX5_XENLA | MASGFSFGTAAASTTTLNPTAAAPFSFGATPAASNTGTTGGLGFGAFN<br>AAATPATTTATTGLGGGLFGAKPAAGFTLGGANTATATTTAASTGFSV<br>GFNKPAGSATPFSLPVTSTSSGGLSLASALTSTPATGPSPFTLNLGST<br>PATTTAAATGLSLGGTLTGLGGSLFQNTNPSATGLGQSTLGQSTLGQS<br>TLGQSLLGQSLLGQSLLGQSTLGQSTLGQSLLGQSLLGLGLNLGAVAP<br>VSQVTTHEGLGGLDFSSSSDKKSDKAGTRPEDSKALKDENLPQLLCQD<br>VENFQKFVKEQKQVQEEISRMSSKAMLKVQEDIKALKQLLSVASSGLQ<br>RNALAIDKLKIETAEELKNAEIALRTQKTPPGLQHENTAPADYFHTLV<br>QQFEVQLQQYRQQIEELENHLATQSNTLHLSPQDLSMAMQKLYQTFVA<br>LAAQLQAVNENFKMLKEQYLGYRKAFLGDSTDVFEARRAEAKKWQNAP<br>RVTTGPTPFSNIPNAAAVAMAATLTQQQQPTTGFGSSSAFGGNTSGSS<br>SFGFGTANKPSGSLSAGFGSTSTSGFNFSNPGINASAGLTFGVSNPSS<br>TSFGTGQLLQLKKPPAGNKRGKR |
| Q9JMD0-3 | ZN207_MOUSE | MGRKKKKQLKPWCWYCNRDFDDEKILIQHQKAKHFKCHICHKKLYTGP<br>GLAIHCMQVHKETIDAVPNAIPGRTDIELEIYGMEGIPEKDMDERRRL<br>LEQKTQESQKKKQQDDSDEYDDDESAASTSFQPQPVQPQQGYIPPMAQ<br>PGLPPVPGAPGMPPGIPPLMPGVPPLMPGMPPVMPGMPPGLHHQRKYT<br>QSFCGENIMMPMGGMMPPGPGIPPLMPGMPPGMPPPVPRPGIPPMTQA<br>QAVSAPGILNRPPAPTAAVPAPQPPVTKPLFPSAGQAQAAVQGPVGTD<br>FKPLNSTPAATTTEPPKPTFPAYTQSTASTTSTTNSTAAKPAASITSK<br>PATLTTTSATSKLIHPDEDISLEERRAQLPKYQRNLPRPGQTPIGNPP<br>VGPIGGMMPPQPGLPQQQAMRPPMPPHGQYGGHHQGMPGYLPGAMPPY<br>GQGPPMVPPYQGGPPRPPMGMRPPVMSQGGRY |
| C3XWA2 | C3XWA2_BRAFL | MFGQQKTPFGGTTGFGTGAFGTSSFGATQTPATGGLFGGTATNTGTGL<br>FGGTSFGTPSTSTSTFGGFGTSTTQTGGGLFGTSTSTAGTGLFATPQQ<br>QTAPFGAANKTGFGGFGTQTSTAATGTGLFGATQQTPSLFGGGQTTST<br>GLFGAVGGIAAGTNGTTVKFNPVSGSDTMMKNGVSQNIRTAHQCITAM<br>KEYETKSLEELRVEDYLANRKGGSTGTTAMFGATATPQTGGGLFGNTA |

| | | |
|---|---|---|
| | | TTTSTTGFTFGKAAFGTGQTQAKGATGFGTTSTGTGLFGQTQTTQAGG<br>LFASPFGGTATTTTPSTGFSFGQTNTGTGLFGQTQQKTGLFGQPTTQT<br>TGLFGTPSTTTTTGFGTTFGTGTFGTQNQAGGLFGANKAPTFGATTTT<br>STTGGLFGNTATNTGGLFGQNKPGLTLGLGTGFGTGAFGTTTTSTGTS<br>LFGPKPTNTFGAGLGTGLGAGLGTLGTGSIFGNTGLGAGIGTGTTGLN<br>LTQPAAALGTDAAAQLAQQAQIQQQLQALSNSPFGDSPLFRNNVADPA<br>KRAELLKPTSLAAQKALTTPSHYKISPRPTAKIKPKPLHSLVAGKSQL<br>FEGLDDEDTSLSNDTFVPRRSVKKLIIRNKASPTASELDAGLGEEDDL<br>LAPPAYPSDTFLVEDKRTSESEKENDPMENLYSNPVRKPIPETPANKS<br>SPAPDDSIIALNVRQRSCSPGQDSMGEPTSDDQGEEEVEREPHPAGII<br>LTKNEYYTEPSLDELANMVDENGDCWVENFVIGREGYGSVFFPGLTNV<br>ANLNLDETVHIRRKEITVYPDDSTKPPEGDGLNKKAEVTLHCTWPMDK<br>TAHIPIKSPDRLKKMGYQEKIEVATSKIGARFLEYRPDTGSWVFQVPH<br>FSKYGLDDSEDEEELVGQDQKKMKMQQQTDLQKQQLQKENQPPSLNKQ<br>QPITERVKSEQPIMEQMQDDSDMADISQEPIPDLAPNSSGDQDGLEGL<br>SEEEVVPSSHRLATHLGMSAHRMQVMKASFFGDEEEGEGMDYKPVFGG<br>MSGRSSLRGSAPPSGAASPLPHRMAESKMDRARLGSLGSPVRGLGGMS<br>PSHSPRTRIPSGGFSPKYSPKVEQSGLFPSLSKPAQYPVFERNQGLLL<br>PSGLSSEIERPRKIVGSQRQPMLLPWRETVTNEKQSLVADAALMMGRS<br>FKVGWGPNWTLVRCAGADREGEAKERKKETSIPFSILPKSTPRLAKLG<br>TSSPFSVVMERVDVAPHFVVQDKVTVANHVSHLEVELETSLCSTEDHP<br>CPVFVPTPGVNALHQHAEVAGRNRENTGVRHPERESADHASLVLSLCV<br>ALWGDLPDQDREDGEDGLDRTSYAYHMARREAVSHWLMNQAQDTIEQE<br>VQDSKFKEGGHTDAIFSLLSGRQISQACSLAQQSGDHRLALLLAQASG<br>SHFPRELVGKQLSDWEELKADRFISDSRLRVYALLAGKPVWPATNGEI<br>NTCENLDWKRALAVHLWYLCTPSATVSEALHLYCKAFTKNNSDYGEYA<br>NPPLPPYLERSADRSGDQEDRYVIRDMCYHLLQLYADRSHQLDRLLAP<br>TTSTPNQLDYRLSWLLCQALQALDYTHLSEHHLNTIHAGYIAQLESLG<br>LWEWAVFVALHITDNSRRELVTKELLCRHCSLSEEEAYVQKEEFLQEK<br>LRVPVVWIHEAKALRAQYEGKSHDEAWHLLKAEKWNASHRIILKHLAA<br>DAIINEDYEYLKEFLEELSPPDRSSTIQDWNFGGRVFLDYVLINNALV<br>EMAKGNSSSYDLERLHPQVTSLCSRIENIPCLTPKDRLCQSEMAKKAA<br>SFLRLVLEGQNKPPTVDGEYLVPSYQLAPHITKLPMPEDYALQELRAL<br>THSYMMELTT |
| J7I6Y1 | J7I6Y1_XENTR | MFNKTFGSPFGTGNGAFGATSTFGQTTGFGTTPATAFGSAGFGTNTST<br>GGLFGNTQTKPGGLFGSTTFNQPATSSSSSGFGFGASTGTTNSLFGST<br>NTGSGLFATQSNAFGQAKPTTFGNFGTSTSTGGLFGNTNTANPFGGTS<br>ASLFGASTFSAAPTGTTIKFNPPSGTDTMAKGGVTTNISTKHQCITAM<br>KEYESKSLEELRLEDYQANRKGPQNPVGAPTGTGLFGTSAATSSASTG<br>IFGSTAANNSFSFAGNKTTFGTAGTGAFGGNTGGLFGQPANQPAASLF<br>NKPFGNATTTQSTGFSFGNTSTLGQPQTSTMGLFGANQPTQSGGLFGT<br>TTNTNATGAFGAGTSLFGQPNPAPFGTGSTLFGNKPAGFGTTTTSAPA<br>FGTTTGGLFGNKPTLTLGTNTNTSNFGFGSNTAGTSLFGNKTATGTIG<br>PSLGTGFGTALNPGQTSLFGSNQPKLTGTLGTGAFGNAGFNSTSAGLG<br>FGAPQAPVAALSDPGASAAHQMFLQQQYNALRYSPFPDSPLFRNPISD<br>PKKKEELLKPTNPAAQKAVLTPTHYKLTPRPAARVRPKALQNSGAAKS<br>QLFDALDDDEPVLSSGVFMPKKSIKKLDLKYLKNSSSLFGQENRDVDD<br>MVPPTPEAPERSVENHHGDEENDEEAATTYPSPFSRPLPQSQEKKQVN<br>HMDDTIVALNMRKSGRIGLEHSSEDASFNEDSFRETEILDASPHPAGI<br>ILTRDSYYTIPSMEELARSVDENGECIVNGFTIGREGFGSIYFEGIVN<br>LTNLDLDSIVHIRRKEVIVYVDDQNKPPLGEGLNRPAQVTLDEVWPID<br>KTSRCMITSPERLSEMNYKSKLENASRKQGAQFVDYRPESGSWVFKVN<br>HFSKYGLQDSDEEDDQLNSAEAKKLKTAPVPPQGKQPPLQQATLPGKV<br>TPPPQSPAVDQLDRVMELDSDMADITQDQDLDSVAEEQDISEEQEPLS<br>ASSHIASSLGINPHALQVMKASLLLEEEDGEIMSRFSSFPSSMDPYPD<br>VRSPRLFPSSHAKRPSSIGLLQSKFSAPSMSRLSETVQGSHSPKIHPA<br>APWSVPAPLAPSFIMPGPAPDTHLRTVGTRRQQELVPLEKSVTHGRGT<br>LMIDMGLFMGRSFRVGWGPNWTLVHNGDKLSERLNAEEDRDMDTIDYG<br>FLPKPTSAKSLTESPFKVHVEKLSLEQKTKDLQSYLLPLEIELKNSTV<br>DKSGPCPHFRPNPGVTAIHDYAGWVRNFSSEAAEVEAVVKQWGLTWTL<br>CESLWGQLKELEASLDEPNEYVKNLERRKAFSHWLAQTAQERIEEEVS<br>LYGPERHIEAVFSYLTGGRISDACRLAQKSGDHRLSLLLSQMVGSQEV<br>RDLITLQLVDWNKLQVDHYIQEERLRVFCLLSGTPVWRSSDNRSINVC<br>SQLDWKRTLGIHLWYMLPPTATVAQALHMYEQAFQEQEGGEPYACYPL<br>PPYLEDCGFSFGDDPSAKFCSLQRDVCVHLLKLYSERQYDLCQLLDPS |

| | | |
|---|---|---|
| | | SATPDPLDYRLSWHMWMVLQALNYTHLSGHRQGMLHASYAAQLENVGL<br>WEWAIFVLLHIQDPHVREAAVRELLNRHCVVHDSPESLAKENFLIQRL<br>CLPAQWIHKAKAVRSRRDGDKHKEALYLLKSHQWNQCHKLVTRHLAAD<br>AVINENYRYLRGFLGELARPEHCKHIQDWETAGKVYLDYISVIEMLNQ<br>IRQDECSGGELEKLHTKVMSLCKWVELIQCYSAKGRLAQSEMAKRVAN<br>ILRVVLSLQQPPESMSDSSSEPRVPLRLLAPHIGRLPMPEDYALEELR<br>GLTQSYLRELICGS |
| Q9VCH5 | NUP98_DROME | MFGGAKPSFGATPAATSFGGFSGTTTTTPFGQSAFGKPAAPAFGNTST<br>FAAQPAQQSLFGAAATPAQPAGGLFGANTSTGFGSTATAQPTAFGAFS<br>QPQQTSNIFGSTQTAASTSLFGQSTLPAFGAAKPTMTAFGQTAAAQPT<br>GSLFGQPAAATSTTGFGGFGTSAPTTTNVFGSGTASAFAQPQATAVGA<br>SGVNTGTAVAKYQPTIGTDTLMKSGQANSVNTKQHCITAMKEFEGKSL<br>EELRLEDYMCGRKGPQAGNAPGAFGFGAQVTQPAQPASGGLFGSTAQP<br>STGLFGQTVTENKSMFGTTAFGQQPATNNAFGAATQQNNFLQKPFGAT<br>TTTPFAAPAADASNPFGAKPAFGQGGSLFGQAPATSAAPAFGQTNTGF<br>GGFGTTAGATQQSTLFGATPAADPNKSAFGLGTAASAATTGFGFGAPA<br>TSTAGGGLFGNKPATSFAAPTFGATSTASTPFSNFGLNTSTAATGGGL<br>FNSGLNKPATSGFGGFGATSAAPLNFNAGNTGGSLFGNTAKPGGGLFG<br>GGTTTLGGTGAAPTGGLFGGGTTSFGGVGGSLGGGGFGMGTNNSLTGG<br>IMGAQPTLGIMTPSHQPIHQQILARVTSPYGDSPIFKDLKLSSEADAT<br>RATNPAAQQAVLDLTSNQYKISTSNNPAPMKVKALGSTLNRKSLFDGL<br>EEFDASVEGFNLKPSAKRLVIKPKVKSVEGGNPSSSIGSAPNTPQSRP<br>KGATPNKERESFSGAIPSEPLPPAGNSPGATNGRESQDNGRRESWLHP<br>NNLEKVRQHNIQTGMDQGSPHNSTLNELVPRKPLDTYRPSSTVRLSVS<br>TIPENPFEDQSSTIARRETFTSQQANESVLSNRSNEAEDSAANQSRLA<br>IEAAAAEAADDESHPTGIVLRRVGYYTIPSLDDLRSYLAEDGSCVVPN<br>FTVGREGYGNVFFGKEMDVAGLNLDEIVHFRNKEIIIYPDDENKPPIG<br>QGLNRDAQVTLDQVWPLDKTKHEAIKDPQRLLEMDWEGKLRRVCDKND<br>TRFIEYRPETGSWVFRVKHFSKYGLGDSDEEDELPTDPKKAKIATLEA<br>QQRANAEKMTLNSLRQAQKISEDAARNLDPKALVAGVASGFRPMDDTA<br>EFLLMDKTQFFQAGGNSDFSMFDPPRQRPTITSPTAVLAQEMVGNEAH<br>KMQLMKSSFFVEDNAPEDEPMETTGRLLRHRKFFNVEPLVWKDGASES<br>SSQYDFEHPSPALPISSSVSEASLMCDAHYEETSSMATGSIVAAVKET<br>KFEMPVTKAFKFVCKPKVAPIKLRATTVPLPRSIAYEMRDNWIADLGF<br>YKGRSFKLSFGPQNSLVLPSTYNNMQNLKEFTGPSLPVSMVFAPRSAT<br>DLSPSVMQLVEFNMVKGNEGFRESIIPHLEVQLNDCLSVNVEGSECPC<br>IHPDSGTKLVSKHFSESLKQRNAGLKEDYSVSVWSLLFALWGDHDELV<br>DLEKNSHYMVMCRRNLLSEWLENTLLGKDLLSKKVSTHSYLEHMLDLL<br>SCHRVNEACELAFSYDDANLALVLSQLSSGAVFRLLMEEQLFAWQQSK<br>SDKYIDLERLKMYMLAAGAPMMQSSHGAINLLENKNWLTALALQLWYF<br>TAPTSSITDALNAYNDAFQAEECYAEPPKPSYRDAPTDTKKPVYDLRY<br>HLLQLHSKRMHSLEETLNPITHTADAMDFRLSWLLLQTLRALGYRHCS<br>PLTEARLSVDFASQLENEGLWQWGIFVLLHIKQQTQRERAVQQMLQRN<br>VSVSAKVALYAEERFIVEELGIPMSWVDYAKAVKAGASGKHHLQAKYL<br>LKAKHFATAHDVIFQHIAPDAIINGKMKYLHSLLIQFEDTEGSSIRVP<br>NWANQGQIFLDFIDISAKFKQIRSVTNIADINARWENLKPQLSELCSR<br>ISLLPCPTSKHRLCQSEISQSLSCLVHGMCIVCPEMESSTVLKVALER<br>LPLPQEFASKELRIWLEELLDKIQNEPPFSERQQPTMMEI |
| K9ZRR1 | K9ZRR1_XENTR | MAAAGGGGGAGESGTGGKIRSRRYHLSSGRTPYSKSRQRQGIISRVTD<br>AVKSIVPGWLQNYFNKQEEERGRAHNASELIVEDTEGRQNNTEHHIYV<br>DDEGNSASGRVTPEPIINVDEEVPSTSQSAINDTDALTRPSLHRANLN<br>FNLFDSPAMNCQPTTSTFPIGTSGFSLVKEIKDSTSQHDDDNISTTSG<br>FSSRASDKDMAVSKNVGVPPLWSPEVDRSQSLSHNSSMTSKKPTFNLS<br>AFSSLSPSLGNASFVNSQLGDSPFYPGKTTYQGAAAVRSSRVRSNPYQ<br>APLRRQVKAKPANAQQYGVTSSTARRILQSLEKMSSPLADAKRIPSNS<br>LSHTPEKSLMDIAENPSKRKKVESPFPPVQKLVTPKSISVSTNRSLYI<br>KPSLTPSAVSNTSSRRVQPDKHNESRENNVQTVQSTSHSFSYPKFSTP<br>TSNELLSRGGGKMMRQKGSHYSTKPAEETSGLTSSSPLFTFSSPIVKS<br>TESNAQSPGSSVDFTFSVPTVKVSSATTSIDSKVSAVSTTAKTHIAVN<br>NSSAKDSDEQVGFCKPAKTLKEGSVLDILRSPGFSSSPSQQTSASTPN<br>RSTSTFSKTVGNAFSPVKVTLGVGGKQALGLWQCSACCHENMASDSNC<br>TACCAAKPQPAETNKKLPASPQHSNTKTTVPLSSVQGLGDKFKLPAGT<br>WDCDTCLVQNKAEVTKCVACETPKPGTGMKANLLLPPATKALNSAPTP<br>SAFVSSSGSITNEEKPEGSWDCTGCRMQNKTENNTCVACKADKPGAIK<br>SLSTAAPSGLLGLLDHFKKPAGSWDCDVCLIQNKPEATKCVACESAKP |

| | | |
|---|---|---|
| | | GTKPELKGTFGTVQTSAAPVSLGLGLLDQFKKSAGSWDCEVCLVENKP<br>EANKCIACESAKPGTKAELKGFGTSTLSAGTTAPTFKFGVQPSDSAGE<br>LKSGASTDSTSGFSFAKPIGDFKFGLASASATTEETGKKSFTFGTSTS<br>NQASAGFKFGVASSAQTNQDTSGGFTFGSVSSTVSFSPAATYSGSTGL<br>QVPAADDDSSRASAAGLKSAEEKKPEAPAVTAFSFGKTDQNKETVSTS<br>FIFGKKDEKTDSAPTGNSFGFGLKKDGEEPKQFLFGKPEPTKEDSTSS<br>TASAGFAFRVSNPTEKKDVEQPVKSVFAFGSQTSTTDAGASKQPFSFL<br>TGVSSTSASSSASGVSSSVFGSVAQSSTPANPSNVFGSATSSNPPAVS<br>SGVFGNLNPSNAPASSSTLFGNVAPSSTPSGSSSLFGTANPSSTPASS<br>SSLFGTAAKLSAPVGSGGVFNSAAPVPPASTSSSVFGSAAPANTSANS<br>ANIFGSAGGTSGAPGTFVFGQPASTTSTVFGNSSESKSTFAFSGQETK<br>PVTSAITSATPFVFGAESASTTPAAPGFNFGRTNTSNVTGTSSSPFIF<br>GGGPTASASPSLTAHANPVPAFGQSANSSTAPAFGSSTSVFPAGNSQQ<br>VPAFGSSTAQPPVFGQQAAQPSFGSSAAPSAGSGFQFGNNTNFNFTTP<br>NSSGGVFTFGANAGSTPQPPAPGFMFNAAASGFNVGTNGRSTPASSIS<br>NRKIKTARRRK |
| Q96PK6 | RBM14_HUMAN | MKIFVGNVDGADTTPEELAALFAPYGTVMSCAVMKQFAFVHMRENAGA<br>LRAIEALHGHELRPGRALVVEMSRPRPLNTWKIFVGNVSAACTSQELR<br>SLFERRGRVIECDVVKDYAFVHMEKEADAKAAIAQLNGKEVKGKRINV<br>ELSTKGQKKGPGLAVQSGDKTKKPGAGDTAFPGTGGFSATFDYQQAFG<br>NSTGGFDGQARQPTPPFFGRDRSPLRRSPPRASYVAPLTAQPATYRAQ<br>PSVSLGAAYRAQPSASLGVGYRTQPMTAQAASYRAQPSVSLGAPYRGQ<br>LASPSSQSAAASSLGPYGGAQPSASALSSYGGQAAAASSLNSYGAQGS<br>SLASYGNQPSSYGAQAASSYGVRAAASSYNTQGAASSLGSYGAQAASY<br>GAQSAASSLAYGAQAASYNAQPSASYNAQSAPYAAQQAASYSSQPAAY<br>VAQPATAAAYASQPAAYAAQATTPMAGSYGAQPVVQTQLNSYGAQASM<br>GLSGSYGAQSAAAATGSYGAAAAYGAQPSATLAAPYRTQSSASLAASY<br>AAQQHPQAAASYRGQPGNAYDGAGQPSAAYLSMSQGAVANANSTPPPY<br>ERTRLSPPRASYDDPYKKAVAMSKRYGSDRRLAELSDYRRLSESQLSF<br>RRSPTKSSLDYRRLPDAHSDYARYSGSYNDYLRAAQMHSGYQRRM |
| K9ZTJ6 | K9ZTJ6_XENLA | MAFNFGATTGTPANQGTTGFSLGTFTPKTTTSGFGFGTTTTTAPTGFG<br>GGFGGFGATTTASTGPAFSFTTPANTTSGLFGATQNKGFGFGTGFGST<br>TTSTGLGTGLGTGLGFTGFNTSQQQQQQSVLGAGLFNQSFQSTPQSNQ<br>LINTASALSAPTLLGDERDAILAKWNQLQAFWGTGKGFFMNNTPPVEF<br>TQENPFCRFKAVGFSYIPNNKDEDGLISLIFNKKESDIRGQQQQLVES<br>LHKVLGGHQTLTVNVEGVKTKADNQTEVIIYVVERSPNGTSRRVGASA<br>LFSYFEQAHIKANMQQLGVTGAMAQTELSPVQIKQLIQNPLSGVDPII<br>WEQAKVDNPDPERLIPVPMIGFKELLRRLEVQDQMTKQHQSRLDIISE<br>DIGELQKNQTTTMAKIGQYKRKLMELSHRVLQVLIKQEIQRKSGFAIQ<br>AEEEQLRVQLDTIQSELNAPTQFKGRLNELMSQIRMQNHFGAVRSEEK<br>YYVDADLLREIKQHLKQQQEGVSHLISIIKDNHEDIKLIEQGLNDNLH<br>MRTGFLS |
| Q7ZXV8 | ZN207_XENLA | MGRKKKKQLKPWCWYCNRDFDDEKILIQHQKAKHFKCHICHKKLYTGP<br>GLAIHCMQVHKETIDAVPNAIPGRTDIELEIYGMEGIPEKDMEERRRI<br>LEQKTQVDGQKKKTNQDDSDYDDDDDTAPSTSFQQMQTQQAFMPTMGQ<br>PGIPGLPGAPGMPPGITSLMPAVPPLISGIPHVMAGMHPHGMMSMGGM<br>MHPHRPGIPPMMAGLPPGVPPPGLRPGIPPVTQAQPALSQAVVSRLPV<br>PSTSAPALQSVPKPLFPSAGQAQAHISGPVGTDFKPLNNIPATTAEHP<br>KPTFPAYTQSTMSTTSTTNSTASKPSTSITSKPATLTTTSATSKLVHP<br>DEDISLEEKRAQLPKYQRNLPRPGQAPISNMGSTAVGPLGAMMAPRPG<br>LPPQQHGMRHPLPPHGQYGAPLQGMAGYHPGTMPPFGQGPPMVPPFQG<br>GPPRPLMGIRPPVMSQGGRY |
| Q91349 | Q91349_XENLA | MSGFNFGAASAGGFSFGNPKSTTTTAPTGFSFGAATAAPSGGFSFGTA<br>TPTPASTTGQTSGLFSFSNPAPSLAPTSGFSFGAQVTSTPAPSSGGLA<br>FGANTSKLNSGVGNQPAGGTTQTSQPMGGFSFGAATTQTQPSATSVGG<br>FSFAGGVGSTSTNVFAQPAASTGITLQSAVSTAAAPTATTSQPTSTFS<br>FGTQPQAAPALNFGLLSSSSVLSTASTPAAAQPVAPTTGLSLNFGKPA<br>DTSAAVTSTGSTTTNTPSLSSLLGTSGPSLFSSVATSTVPSVVSTVAS<br>GLSLTSTATSTGFGMKTLASSAVPTGTLATSTASLGVKAPLAGTIVQA<br>NAVGSAAATGISTATAMTYAQLENLINKWSLELEDQEKHFLQQATQVN<br>AWDRTLMQNGERITTLHREMEKVKLDQKRLDQELDFILSQQKELEDLL<br>TPLEESVKEQSGTIYLQHADEEREKTYKLAENIDAQLKRMAQDLKEVI<br>EHLNTSAGPGDASNPLQQICKILNAHMDSLQWIDQNSALLQRKVEQVT<br>KECESRRKEQERGFSIAFD |

| | | |
|---|---|---|
| Q5EWX9 | Q5EWX9_XENLA | MEGGCCWDLVKRIQPIALPLALSLVTCALAYIYNQLTVAGLLLASCWT<br>YRSSGRRFIVWKWLTGPRLVASTKKLRNSTDWRLQSPPVSLLEGSPSP<br>LSLNMGTYMNKQELPARGEACGPRAIKEKLSRPNPSVATPIRRLSFRD<br>PLSSSNRTYLCARRDYPLKQEIYSKPGSLPRVSLDGQSQQRVPLTPRH<br>FTRRTPVKISPQDLKTRFQLAPQPNVVSSPHLDFEQEPCLPSIYHRVE<br>HETSVTHSRHLLEPEQYPASPSSHKANTFEATFSSHHVEDPCATESVL<br>RALKECLKRKRIRDEDSEDTENKRRKENGEESKLLVNDSLCRTSKMEA<br>SRASWRSESSSMENLSQQSSISIQDMLNEVQNSKNSDGKVSPASSFNL<br>ENLSKRKIITVPGHNEITSSLSSSRYLQKKKRSLEIPRCDTPEWPIKK<br>MRKDRLETSMSPSQETMKTQMSETQKVEVNPSPVQLHKPKGNLPWKRN<br>VHLICPEHPEEFYRLPPGPIPGYNVTQADYDAAKEAAQQRFLRLFQEP<br>SAPSVPSASESCSKQVSLTLQTSTSPDKSFQAVNEKPNISAANTTFTL<br>FSSSSDAKSVTSSSSTPSPFTSATTSITNTLLQSLGSGQTKSESPFPK<br>NSLLQILGKTEGNNSQPMFNPVFGPLGSANPSPVTAAPGLSTTTTAIL<br>KPICGDPPSQQAQTSMFKPIFEPPSSQTVPSALSSPFVFSSSSSTTST<br>TSFLGTTGNSVNTGKHEKPNLALTTCASNTTITSSLAPTFGITSKVEP<br>VSLGGPASQANTSFSVIGSTSVPAISTAFGSTTSAFTAAGQTANPSFS<br>ANTAPIGVSSVSASEGQTSISTTATFSKFGVPVGQNVFSASNPFGSGT<br>QSTMGTSTQSATNIAFSFGTSTSTQSAFGFSQNPTMLFSSSTKSNNPT<br>NTSGFNSMGSVFGSSTVPTSSIVTPNKSLSTLGIPEKCETKNQLVTNQ<br>LSLGQGSTPAPFPSIMPTQSFVSSSTSFAPSTPTVNQSSTPGSFPSMA<br>AVQPFTSPSSPAAGFFSHGTAPKSRTAVRHKLHPRRPHPRKK |
| P34761 | WHI3_YEAST | MQSSVYFDQTGSFASSSDNVVSSTTNTHNISPSHRSSLNLNTTSHPHE<br>ASGRGSASGELYLNDTNSPLAISSMLNTLALGSMPQDIASSNISNHDN<br>NIKGSYSLKLSNVAKDITLRECYAIFALAEGVKSIELQKKNSSSSITS<br>ASLEDENDIFIIARFELLNLAINYAVILNSKNELFGPSFPNKTTVEII<br>DDTTKNLVSFPSSAIFNDTSRLNKSNSGMKRPSLLSQRSRFSFSDPFS<br>NDSPLSQQQSQQQQQQPQQPQQHSTQKHSPQQCNQQQVNSSIPLSSQG<br>QVIGLHSNHSHQDLSVESTIQTSDIGKSFLLRDNTEINEKIWGTSGIP<br>SSINGYMSTPQPSTPTLEWGNTSASQHGSSFFLPSAASTAIAPTNSNT<br>SANANASSNNGASNNGANQALSASSQQPMMQIGNTINTSLTSSNSLPP<br>YGLMSSQSQHISNMVNTSDMNITPQKQNRFMQQPQPEHMYPVNQSNTP<br>QKVPPARLSSSRNSHKNNSTTSLSSNITGSASISQADLSLLARIPPPA<br>NPADQNPPCNTLYVGNLPSDATEQELRQLFSGQEGFRRLSFRNKNTTS<br>NGHSHGPMCFVEFDDVSFATRALAELYGRQLPRSTVSSKGGIRLSFSK<br>NPLGVRGPNSRRGGSGNPNPNVNMLSSYNSNVGHIKN |
| Q5XGN1 | NUP42_XENLA | MAICNFFLQGRCRYGEKCWNEHPRGGGGGGGNRYQSQNRYQEQSRYQE<br>QSRYPEQSRYPEQNRYQEPAGNAKGTWGASSQRYVQPSNFSKSTTWIN<br>RDSEKPSAGSFSGFGSRNVKSTAATGLPSTQNRFAALSSQDNSRDGQT<br>DKGNILDDIMKDMEIWESSGQWMFSVYSMLKEKKNISGFTDFSPEELR<br>LEYSVCQAEGNPLKYINAVQQLGSKWKQRILELKNPNPSIKTALLNEL<br>NSPSPDVTPGYSGQQNSAFGALSFPTSNTAPTAVTFSFKADTTTAAKP<br>AVPNALAGSDFSAFGNKPTSAPSFGSGVAAAAASFSFAPSTISGFGST<br>ASNSGFGAASNAAGFQGAANIAAAPAFGVASSTAPASGFGGGFGTTVN<br>TGAKTSSVRDLFSAGTAVPVQTTLLFGQATGSLNTTASSTSLAGQPFK<br>ASTSATAVSGSFTSDNTSNPLFTPRNELSVEDLAQFEAKQFTLGKTPI<br>KPPSADLLKVT |
| Q6DEC7 | Q6DEC7_XENLA | MAKRIADKELTDRNWDQEEEEEDAGMFSVASQEVLKTRAIKKAKRRNA<br>GNESESGGAFKGFKGLFLVTGGSLSSFGNGSPAKPSEGLSNGTSTSLF<br>INLKPQSKPTFGSAFTNRPLLGTAEKSTNGEKPLSSSGAALSKPGNLE<br>YNKQLTSLNCSVRDWIVKHVNANPLCDLTPIFKDYEKHLSAIEQKYGA<br>SSESGSESDGAAQTKTIPNLSSGKTVSIATFSFGNKDKAPEAPTKTPP<br>DSKPQAAPTFNFGQKVDSSTLGLISSGGAPNFSFSIGAPSLFGKNNGS<br>TASSSSSQESEPSGKTEEKGEGEEEEEPPKEVIQEIKEDDAFYSKKCK<br>LFYKKDNEFKEKGVGTLHLKPVENKKTQLLVRADTNLGNILLNILVQP<br>SMPCSRTGKNNVMIVCVPNPPVDEKNPTVPVTLLVRVKSAEDADQLHK<br>ILLEKKEV |
| Q9TXM1 | Q9TXM1_CAEEL | MSSSKPYPSGLPNSRRKRGGRRSSSRSNQESASNNMEHQITLDELFNP<br>IAKQDSAQSTSREYGAKSGISHHGSVSFNGNTFMNGQQLNHSMTRHGR<br>VFNQSMHAAQGNGSNAFNSIPPTAPVFSADFRRNLQTRNSSSWYERRF<br>PVSTDQDDVQQSNTRRSRSRQNGQHGLSFSDGSNNYGHAGNKSFSVSS<br>VPVGFQKQENNSKKLRQTNVHQQCLGNKSFNAQAGVHGHAFKKGHKDN<br>KNASGKEVINSSLVQKHDAIKSRNLNQSFSGFPTHETSSMKNQQQKSR<br>NDRKKSRGSSNFQDRTYFNTNDDELTDDVFIDDSMDAARGRRSRSVTK |

| | | |
|---|---|---|
| | | KLQQSTYSKQNAGSKQLTEKCKSSEEAAKRNLVSNVFSKDGTELSIEQ<br>LLEIVSMKIGQQIHLPSSSHGECSNLNRTLPASDLNCSIGEDFDSSFV<br>DANNQTLPVSLPKKTSLSIKRRGSSRSASRLASLDVTLETVEEDEEPT<br>PSPQPSSPPKISRRKWTGTFDANVEEMRRLLHGDPEMPKSANRASSSK<br>DQINRNNVDVKRTPSSSIIPTPKALIGERCLTSSSKSSKLNKSLGVVD<br>SKATKSPMYSVTVSGKETASGKRIAQKLTPKVVALESSYITGIPVSTD<br>CNGCPTPKRSGINCEIRAAEVYNQAGKWPFEITSDPAPLPCESADRIE<br>YPSQDCTQDPASTSPPPRISESLTAFLEAQQDFNDYIDTNYKEKTQLL<br>KVNLNIHGMSPERWLYLNYFCTETIPRLDGPYADDPRVPPVRNMFRKW<br>FLRFAEACLGNPHQLAVMQEIAATFVQARLDDTSSSTDSTNMLYMLWK<br>ECIGQKNIIAIADACLLAHLRKSDPIKYLNVKRDWLESIFDPPRDQ |
| P39936 | IF4F2_YEAST | MTDQRGPPPPHPQQANGYKKFPPHDNQYSGANNSQPNNHYNENLYSAR<br>EPHNNKQYQSKNGKYGTNKYNNRNNSQGNAQYYNNRFNNGYRLNNNDY<br>NPAMLPGMQWPANYYAPQMYYIPQQMVPVASPPYTHQPLNTNPEPPST<br>PKTTKIEITTKTGERLNLKKFHEEKKASKGEEKNDGVEQKSKSGTPFE<br>KEATPVLPANEAVKDTLTETSNEKSTSEAENTKRLFLEQVRLRKAAME<br>RKKNGLISETEKKQETSNHDNTDTTKPNSVIESEPIKEAPKPTGEANE<br>VVIDGKSGASVKTPQHVTGSVTKSVTFNEPENESSSQDVDELVKDDDT<br>TEISDTTGGKTVNKSDDETINSVITTEENTVKETEPSTSDIEMPTVSQ<br>LLETLGKAQPISDIYEFAYPENVERPDIKYKKPSVKYTYGPTFLLQFK<br>DKLKFRPDPAWVEAVSSKIVIPPHIARNKPKDSGRFGGDFRSPSMRGM<br>DHTSSSRVSSKRRSKRMGDDRRSNRGYTSRKDREKAAEKAEEQAPKEE<br>IAPLVPSANRWIPKSRVKKTEKKLAPDGKTELFDKEEVERKMKSLLNK<br>LTLEMFDSISSEILDIANQSKWEDDGETLKIVIEQIFHKACDEPHWSS<br>MYAQLCGKVVKDLDPNIKDKENEGKNGPKLVLHYLVARCHEEFEKGWA<br>DKLPAGEDGNPLEPEMMSDEYYIAAAAKRRGLGLVRFIGYLYCLNLLT<br>GKMMFECFRRLMKDLNNDPSEETLESVIELLNTVGEQFEHDKFVTPQA<br>TLEGSVLLDNLFMLLQHIIDGGTISNRIKFKLIDVKELREIKHWNSAK<br>KDAGPKTIQQIHQEEEQLRQKKNSQRSNSRFNNHNQSNSNRYSSNRRN<br>MQNTQRDSFASTKTGSFRNNQRNARKVEEVSQAPRANMFDALMNNDGD<br>SD |

[a] This protein set was curated by Vernon *et al.* (1).

### Table S2. Summary table for the set of proteins that exhibit homotypic phase separation behavior.

| Seq. # | Seq. length | Σ class. dist. P | longest PS IDR | first residue longest PS IDR | last residue longest PS IDR | Σ class. dist. D | longest non-PS IDR | first residue longest non-PS IDR | last residue longest non-PS IDR | Σ class. dist. F | longest F-labeled region | first residue longest F-labeled region | last residue longest F-labeled region | UniProtKB | Gene | Protein |
|---|---|---|---|---|---|---|---|---|---|---|---|---|---|---|---|---|
| 1 | 592 | 1768 | 266 | 326 | 591 | 16 | 0 | 0 | 0 | 38 | 27 | 242 | 268 | Q92804 | RBP56 | TATA-b |
| 2 | 320 | 514 | 129 | 191 | 319 | 40 | 23 | 74 | 96 | 97 | 74 | 107 | 180 | P09651-2 | ROA1 | Isoform A |
| 3 | 708 | 952 | 214 | 1 | 214 | 12 | 0 | 0 | 0 | 432 | 332 | 319 | 650 | D0PV95 | DDX3 | ATP-depe |
| 4 | 263 | 1067 | 262 | 1 | 262 | 0 | 0 | 0 | 0 | 0 | 0 | 0 | 0 | H3BNZ4 | H3BNZ4 | FUS RNA |
| 5 | 662 | 461 | 95 | 47 | 141 | 37 | 0 | 0 | 0 | 424 | 329 | 259 | 587 | O00571 | DDX3X | ATP-depe |
| 6 | 724 | 459 | 234 | 23 | 256 | 31 | 0 | 0 | 0 | 537 | 452 | 262 | 713 | Q9NQI0 | DDX4 | Probable |
| 7 | 248 | 205 | 77 | 122 | 198 | 46 | 27 | 194 | 220 | 82 | 74 | 46 | 119 | Q15056 | IF4H | Eukaryoti |
| 8 | 823 | 757 | 309 | 1 | 309 | 148 | 40 | 517 | 556 | 101 | 99 | 628 | 726 | P14907 | NSP1 | Nucleopor |
| 9 | 292 | 614 | 291 | 1 | 291 | 0 | 0 | 0 | 0 | 0 | 0 | 0 | 0 | F8WC90 | F8WC90 | EWS RN |
| 10 | 386 | 102 | 42 | 344 | 385 | 23 | 0 | 0 | 0 | 231 | 151 | 205 | 355 | P31483 | TIA1 | Cytotoxic |
| 11 | 786 | 603 | 50 | 104 | 153 | 202 | 25 | 364 | 388 | 118 | 34 | 1 | 34 | P15502 | ELN | Elastin |
| 12 | 341 | 762 | 157 | 184 | 340 | 42 | 20 | 76 | 95 | 93 | 75 | 1 | 75 | P22626-2 | ROA2 | Isoform A |
| 13 | 323 | 231 | 94 | 1 | 94 | 41 | 0 | 0 | 0 | 212 | 189 | 134 | 322 | P22232 | FBRL | rRNA 2'- |
| 14 | 693 | 330 | 82 | 611 | 692 | 101 | 29 | 549 | 577 | 596 | 465 | 1 | 465 | G5EBV6 | PGL3 | Guanyl-sp |
| 15 | 172 | 235 | 98 | 74 | 171 | 4 | 0 | 0 | 0 | 33 | 64 | 1 | 64 | Q14011 | CIRBP | Cold-indu |
| 16 | 1105 | 2224 | 361 | 1 | 361 | 118 | 0 | 0 | 0 | 303 | 141 | 964 | 1104 | D3KYQ3 | D3KYQ3 | Macronuc |
| 17 | 1553 | 1540 | 640 | 1 | 640 | 122 | 0 | 0 | 0 | 898 | 541 | 1012 | 1552 | Q387F2 | Q387F2 | Nucleopor |
| 18 | 2834 | 4397 | 94 | 2646 | 2739 | 0 | 0 | 0 | 0 | 1029 | 117 | 2531 | 2647 | Q64K55 | Q64K55 | Aciniform |
| 19 | 959 | 2000 | 701 | 1 | 701 | 57 | 43 | 747 | 789 | 232 | 161 | 798 | 958 | Q02629 | NU100 | Nucleopor |
| 20 | 1113 | 2004 | 632 | 164 | 795 | 216 | 41 | 920 | 960 | 241 | 130 | 958 | 1087 | Q02630 | NU116 | Nucleopor |
| 21 | 997 | 1706 | 626 | 1 | 626 | 93 | 0 | 0 | 0 | 164 | 107 | 841 | 947 | F4ID16 | NU98B | Nuclear p |
| 22 | 2053 | 2753 | 504 | 276 | 779 | 170 | 35 | 1171 | 1205 | 1119 | 599 | 1454 | 2052 | Q54EQ8 | NUP98 | Nuclear p |
| 23 | 414 | 402 | 79 | 335 | 413 | 58 | 22 | 1 | 22 | 251 | 96 | 96 | 191 | Q13148 | TADBP | TAR DN |
| 24 | 2037 | 1528 | 269 | 1768 | 2036 | 410 | 53 | 1164 | 1216 | 758 | 407 | 15 | 421 | Q9PVZ2 | Q9PVZ2 | Nucleopor |
| 25 | 1678 | 1399 | 320 | 189 | 508 | 209 | 25 | 1018 | 1042 | 883 | 227 | 1451 | 1677 | G5EEH9 | NUP98 | Nuclear p |
| 26 | 1817 | 1299 | 193 | 1 | 193 | 399 | 61 | 886 | 946 | 1186 | 649 | 1168 | 1816 | P52948 | NUP98 | Nuclear p |
| 27 | 1475 | 1353 | 291 | 1184 | 1474 | 219 | 32 | 1150 | 1181 | 290 | 62 | 716 | 777 | P49790 | NU153 | Nuclear p |
| 28 | 599 | 584 | 97 | 18 | 114 | 66 | 21 | 471 | 491 | 224 | 104 | 375 | 478 | Q5EAX5 | Q5EAX5 | MGC8499 |
| 29 | 464 | 137 | 60 | 404 | 463 | 191 | 57 | 224 | 280 | 99 | 94 | 1 | 94 | Q9JMD0-3 | ZN207 | Isoform 3 |
| 30 | 1834 | 1305 | 243 | 213 | 455 | 532 | 104 | 872 | 975 | 932 | 330 | 1504 | 1833 | C3XWA2 | C3XWA2 | Nuclear p |
| 31 | 1790 | 1214 | 290 | 214 | 503 | 538 | 94 | 873 | 966 | 1090 | 578 | 1212 | 1789 | J7I6Y1 | J7I6Y1 | Nuclear p |
| 32 | 1960 | 1137 | 254 | 344 | 597 | 302 | 46 | 1034 | 1079 | 1227 | 715 | 1245 | 1959 | Q9VCH5 | NUP98 | Nuclear p |
| 33 | 1547 | 1395 | 383 | 1164 | 1546 | 297 | 29 | 1017 | 1045 | 234 | 46 | 766 | 811 | K9ZRR1 | K9ZRR1 | Nup153 |
| 34 | 669 | 464 | 155 | 281 | 435 | 106 | 26 | 138 | 163 | 152 | 145 | 1 | 145 | Q96PK6 | RBM14 | RNA-bind |
| 35 | 535 | 378 | 119 | 14 | 132 | 9 | 0 | 0 | 0 | 313 | 382 | 153 | 534 | K9ZTJ6 | K9ZTJ6 | Nup54 |
| 36 | 452 | 164 | 57 | 395 | 451 | 116 | 44 | 92 | 135 | 114 | 96 | 1 | 96 | Q7ZXV8 | ZN207 | BUB3-int |
| 37 | 547 | 513 | 167 | 1 | 167 | 59 | 0 | 0 | 0 | 150 | 113 | 349 | 461 | Q91349 | Q91349 | IL4I1 prot |
| 38 | 1050 | 744 | 65 | 571 | 635 | 181 | 42 | 431 | 472 | 327 | 101 | 1 | 101 | Q5EWX9 | Q5EWX9 | Nuclear p |
| 39 | 661 | 505 | 69 | 1 | 69 | 26 | 0 | 0 | 0 | 213 | 69 | 143 | 211 | P34761 | WHI3 | Protein W |
| 40 | 491 | 429 | 63 | 87 | 149 | 15 | 0 | 0 | 0 | 114 | 90 | 154 | 243 | Q5XGN1 | NUP42 | Nucleopor |
| 41 | 440 | 169 | 32 | 113 | 144 | 272 | 35 | 1 | 35 | 171 | 61 | 354 | 414 | Q6DEC7 | Q6DEC7 | LOC5036 |
| 42 | 862 | 269 | 62 | 140 | 201 | 256 | 46 | 328 | 373 | 317 | 166 | 696 | 861 | Q9TXM1 | Q9TXM1 | Uncharact |
| 43 | 914 | 188 | 79 | 17 | 95 | 572 | 87 | 147 | 233 | 413 | 132 | 694 | 825 | P39936 | IF4F2 | Eukaryoti |

**Supporting Figures**

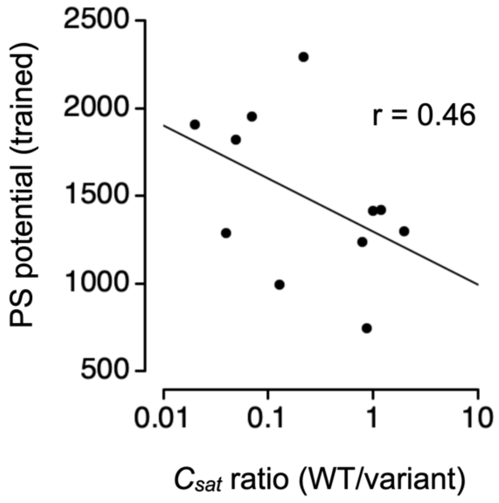


**Figure S1. Mutation effects on experimental $c_{sat}$ compared to the expanded PS potential trained previously using experimental $\Delta h°$ from a mutant dataset.** Experimental $c_{sat}$ ratio (x-axis) was digitally extracted from Figure S18A in Rekhi *et al.* (2); the sequence-calculated PS potential includes the summed P classifier distance plus $U_\pi$ and $U_q$ trained previously (3).
